# Kilohertz volumetric imaging of in-vivo dynamics using squeezed light field microscopy

**DOI:** 10.1101/2024.03.23.586416

**Authors:** Zhaoqiang Wang, Ruixuan Zhao, Daniel A Wagenaar, Diego Espino, Liron Sheintuch, Ohr Benshlomo, Wenjun Kang, Calvin K. Lee, William C. Schmidt, Aryan Pammar, Enbo Zhu, Jing Wang, Gerard C.L. Wong, Rongguang Liang, Peyman Golshani, Tzung K. Hsiai, Liang Gao

## Abstract

Volumetric functional imaging of transient cellular signaling and motion dynamics poses a significant challenge to current microscopy techniques, primarily due to limitations in hardware bandwidth and the restricted photon budget within short exposure times. In response to this challenge, we present squeezed light field microscopy (SLIM), a computational imaging method that enables rapid detection of high-resolution three-dimensional (3D) light signals using only a single, low-format camera sensor area. SLIM pushes the boundaries of 3D optical microscopy, achieving over one thousand volumes per second across a large field of view of 550 μm in diameter and 300 μm in depth with a spatial resolution of 3.6 μm laterally and 6 μm axially. Using SLIM, we demonstrated blood cell velocimetry across the embryonic zebrafish brain and in a free-moving tail exhibiting high-frequency swinging motion. The millisecond temporal resolution also enables accurate voltage imaging of neural membrane potentials in the leech ganglion, and in the hippocampus of behaving mice. These results collectively establish SLIM as a versatile and robust imaging tool for high-speed microscopy applications.

## Introduction

High-speed fluorescence microscopy has been playing an indispensable role in revealing the dynamic interplay and functionality among cells in their native environment. With continuous improvements in fluorescent markers, many transient biological processes, such as the blood flow^1^ and neural action potentials^2–4^, become trackable and, thus, demand microscopy with an ever higher spatiotemporal resolution. Traditional three-dimensional (3D) imaging tools, such as confocal microscopy, light sheet microscopy, and two-photon microscopy, heavily rely on scanning to acquire a volumetric image. Despite advancement in beam shaping^5,6^, remote refocusing mechanisms^7^, detector array^8,9^ and detection geometry^10^, there persists an inherent trade-off between temporal resolution, the 3D field-of-view (FOV), and spatial resolution in these techniques. This constraint marks a significant challenge to obtain optimal performance across a large 3D field of view (FOV) for robust ultrafast detection exceeding kilohertz (kHz).

Computational imaging mitigates this trade-off by encoding high-dimensional information, such as depth^11^, time^12^, and spectra^13,14^, into two-dimensional (2D) multiplexed camera measurements. Among these techniques, light-field microscopy (LFM) excelled in various biological applications, including observation of neural activity in freely moving animals^15–17^ and visualization of hemodynamics in the brain^18^ and heart^19–21^. By simultaneously collecting the spatial and angular information of light rays, LFM enables volumetric reconstruction *post hoc* from snapshot measurements. Without scanning, the sensor bandwidth becomes the primary bottleneck for LFM 3D imaging speed. While modern scientific Complementary Metal-Oxide Semiconductor (sCMOS) sensors typically offer a full framerate lower than 100 Hz, increasing the imaging speed can be achieved by reading out only selected low-format regions of interest (ROI). However, this approach comes at the cost of sacrificing either the spatial and/or angular components associated with the field of view (FOV) and axial resolution.

The integration of ultra-high-speed cameras^7,10,22^ and event cameras^23^ holds promise for providing higher bandwidths to LFM. However, their current limitations in sensitivity and noise performance present challenges, especially for photon-starved applications like imaging genetically encoded voltage indicators (GEVIs) (See **Supplementary Table 1** for a list of camera models and their performance)^7^. On the other hand, the compressibility of four-dimensional (4D) (two spatial dimensions plus two angular dimensions) light fields has been leveraged for compressive detection. Coded masks^24–26^ and random diffusers^27–30^ are employed to modulate and integrate the spatio-angular components originally recorded by distinct pixels. Sparse nonlocal measurements can also be utilized across different angular views to acquire light fields with sensors of arbitrary formats^31–33^. Nevertheless, as compressive imaging relies on sparsity priors and optimization algorithms for signal recovery from the sub-Nyquist measurement, the performance is prone to degradation in challenging scenarios. These methods are primarily validated on photographic scenes and biological samples with relatively long exposure time. Their robustness and effectiveness in kilohertz microscopy with extremely low photon budget such as voltage imaging remain elusive. To address the unmet need for high-speed light-field imaging, we present herein Squeezed LIght field Microscopy (SLIM), which allows the capture of 3D fluorescent signals at kilohertz volume rates in a highly data-efficient manner. SLIM operates by acquiring an array of rotated 2D sub-aperture images. An anamorphic relay system applies anisotropic scaling, effectively ‘squeezing’ the image along one spatial axis. This allows the camera sensor to detect the light field using only a low-format letterbox-shaped ROI. Leveraging the row-by-row readout architecture of modern CMOS sensors, SLIM achieves a more than fivefold increase in acquisition rate compared to traditional LFM. Each squeezed sub-aperture image complements the others, facilitating high-fidelity, robust 3D reconstruction from compressed measurement. By calculating the product of space-bandwidth product (SBP) and volume rate, SLIM measures ∼7.3 gigavoxels per second, making it one of the fastest 3D fluorescent microscopes in literature (**Supplementary Table 2**). We demonstrated SLIM by capturing the flowing red blood cells in free swinging tails of embryonic zebrafish at 1,000 volumes per second (vps), *ex vivo* voltage imaging in dissected leech ganglia at 800 vps and *in vivo* voltage imaging in hippocampus from behaving mice at 800 vps. SLIM enables tracking high-speed cellular motion across a 550 μm FOV within a 300 μm depth range. It allows detection of millisecond membrane action potentials and subthreshold oscillations in 3D space over extended time periods in awake, free-behaving animals. Furthermore, we showcased that the high framerate of SLIM could be exploited to enhance axial resolution when combined with multi-layer scanning light-sheet microscopy. This allows for imaging densely labeled structures, previously challenging with LFM, such as contracting myocardium in a zebrafish, at 4,800 frames per second (fps), leading to a volume rate of 300 vps.

## Results

### Principle and design of SLIM

In a typical SLIM camera (**Fig. 1a**), the input scene is imaged by a combined system consisting of an array of dove prisms and lenslets (**Fig. 1ai**, **Supplementary Figure 1**). Each dove prism within the array is rotated at a distinct angle relative to its optical axis. This arrangement gives rise to an array of perspective images, each rotated at twice the angle of its corresponding dove prism’s rotation, all converging at an intermediate image plane situated behind the lenslets. Subsequently, these rotated perspective images are further processed through an anamorphic relay system consisting of two cylindrical doublets with orthogonal optical axes (**Fig. 1aii**, **Supplementary Figures 2**). This relay system imparts anisotropic scaling to the image array, where the images experience de-magnification (0.2X) along one spatial axis while preserving the original magnification along the orthogonal direction. Finally, the rescaled image array is acquired by a 2D camera, where we read out only pixel rows that receive light signals (referred to as active readout ROI in **Fig. 1a**).

**Fig. 1.**
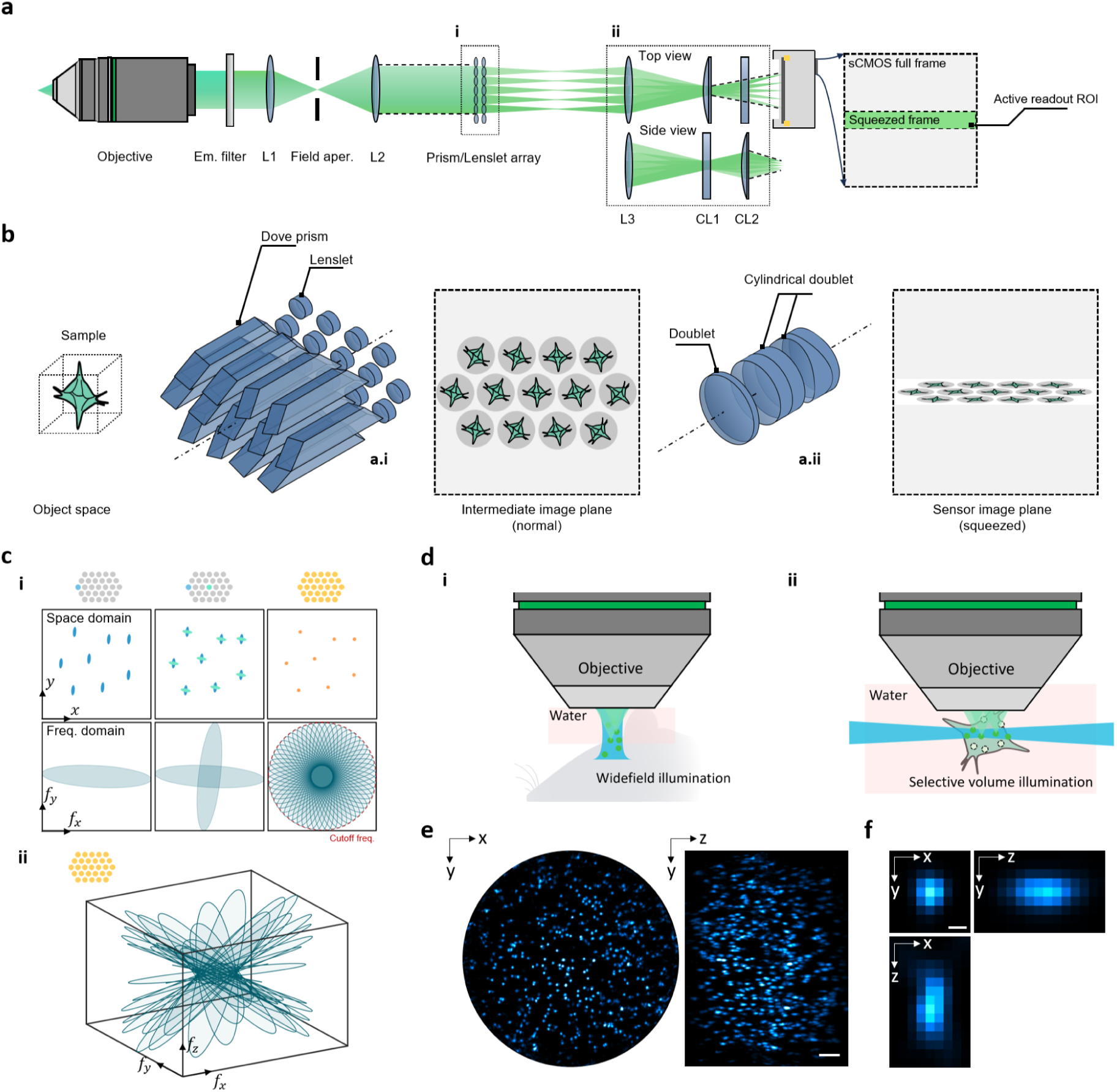
Principle of SLIM. **a.** Schematic of SLIM detection system. L1-L3, achromatic doublet. CL1-CL2, achromatic cylindrical doublet. SLIM records light field at kilohertz frame rates by using a reduced active readout ROI on the camera. **b.** The optical transformation in SLIM comprises image rotation and squeezing, performed by a dove prism/lenslet array (**a.i**) and a customized anamorphic relay (**a.ii**). **c.** (i) Each sub-aperture, after reversing the squeezing and rotation, gives an image with anisotropic spatial resolution but complementary to others. By merging different sub-aperture images, SLIM estimates the original features. Three columns show how one, two and all sub-aperture images reconstruct the final image and spectrum. The red dotted circle denotes the lateral cutoff frequency. (ii) 3D illustration of transfer functions of SLIM by analyzing all sub-aperture images as geometrical projections. Each gives an elliptical slice in the 3D frequency space, depending on its rotation angle and sub-aperture location. **d.** SLIM is compatible with different illumination modes. We demonstrated widefield illumination (i) for behaving mice imaging, and selective volume illumination (ii) for zebrafish and leech ganglion imaging. **e.** 3D MIPs of fluorescent beads. Scale bar, 50 μm. **f.** Cross-sections of single bead showing representative PSFs. Scale bar, 3 μm.

One of the key advantages of employing squeezed optical mapping is the improved readout speed. Modern CMOS sensors are equipped with parallel analog-to-digital converters (ADCs) for each column of pixels, ensuring consistent frame rates regardless of the number of pixel columns being readout. The frame rate is, therefore, solely determined by and inversely proportional to the number of pixel rows being readout^34^. For example, on the Kinetix sCMOS developed by Teledyne, using an ROI of 200×3200 pixels allows SLIM to capture a 19 sub-aperture image array at 1,326 fps and 7,476 fps in 16-bit and 8-bit mode, respectively. In contrast, the full-frame mode achieves frame rates of only 83 fps (16-bit) and 500 fps (8-bit).

The forward model of SLIM is illustrated in **Fig. 1b**. Similar to Fourier LFM (FLFM)^16,35–37^, SLIM can be conceptualized as a tomographic system, where each sub-aperture image is essentially a parallel projection of a 3D volume along a line of sight at the sub-aperture’s view angle^38,39^. However, unlike FLFM, where these sub-apertures images are directly captured by a 2D camera, SLIM applies in-plane rotation and vertical scaling operations to these images before recording. In the 3D spatial frequency space, the Fourier spectrum of a SLIM sub-aperture image manifests as a 2D elliptical slice (Fourier Slice Theorem, **Supplementary Note 1**). The short axis of this ellipse corresponds to the low-resolution sampling along the squeezing direction. By using an array of sub-aperture images rotated at complementary angles, SLIM fills in the missing high-frequency information. This process results in a synthesized power spectrum with a bandwidth that approximates that of the original unsqueezed FLFM (**Fig. 1c**). In addition, the rotation angles of sub-aperture images are carefully crafted to maximize the horizontal projections of their 3D point spread functions (PSFs) (**Supplementary Figure 3**). In other words, when imaging a 3D object, the sub-aperture images of SLIM exhibit lateral disparity shifts due to their view angle difference. While the entire set of rotation angles are sampled uniformly from 0 to 180 degrees, we optimize the rotation angle assignment to each sub-aperture image to align its disparity shift with the unsqueezed spatial axis (*i.e.*, camera pixel row direction), thereby maximizing the samplings of disparity and consequently enhancing the axial resolution (**Supplementary Note 1**). With this approach, some sub-apertures may not receive their optimal angles. We prioritized the sub-apertures in the outer region, as they play a key role in determining the system’s axial resolution. This forward model can be further extended to wave optics by using a sum of 2D convolutions between the sample sliced at each depth and the corresponding sub-aperture PSF^39^. Through an iterative deconvolution algorithm, SLIM reconstructs the 3D fluorescence distribution by fusing all sub-aperture images (**Methods**).

We built two setups to demonstrate SLIM’s applications across a wide range of high-speed biological processes (**Fig. 1d**, **Supplementary Figures 4 and 5**). A widefield epi-illumination setup (**Fig. 1di**) is used for mouse imaging through cranial windows, while a selective volume illumination setup (**Fig. 1dii**) is designed for small animals like zebrafish larvae and dissected leech ganglia, where the target of interest can be accessed via side illumination. Selective illumination, implemented with either a scanning light sheet or slit-confined LED (**Supplementary Figure 4a**), suppresses fluorescence outside the imaging volume, particularly in scattering tissues (**Supplementary Figures 6 and 7**). The LED is preferred for applications requiring high power stability.

**Figure 1e-f** show the 3D reconstruction of fluorescent beads of sub-diffraction size imaged by SLIM. At a magnification of 3.6X, the imaging volume spans a 3D space of Ø550 μm × 300 μm (Ø denotes diameter) with a spatial resolution of 3.6 μm laterally and 6.0 μm axially (**Supplementary Figures 3**). SLIM inherits the lenslet array-based optical design of FLFM for snapshot 3D imaging and provides a highly efficient data acquisition strategy for modern scientific camera sensors. SLIM can record and reconstruct 7.3×10^9^ effective voxels per second, making it a powerful optical compression method that compresses kilohertz volume rates into the limited bandwidth of the camera.

### Imaging of flowing red blood cells in an embryonic zebrafish

Experimental characterization of blood flow in living organisms provides valuable insights into local metabolism, vascular development, and disease states. Using fluorescently labeled blood cells, various imaging methods have been demonstrated in single-cell velocimetry, such as in the larval zebrafish heart^10,20,40^, tail^41,42^, and mouse brain^1,18^. However, these methods are often limited to 2D imaging or restricted by a limited volumetric frame rate, which hinders the detection of fast flow and necessitates sedation of the animal to reduce motion artifacts. Here, we show that SLIM could be used to capture the fast-circulating red blood cells (RBCs) in a zebrafish at a kilohertz volumetric rate, both with and without sedation.

We imaged the transgenic zebrafish embryos expressing DsRed in RBCs at three days post fertilization (dpf). We excited the zebrafish brain using light-sheet-synthesized volumetric illumination and recorded fluorescence using SLIM with 19 sub-aperture images at 1,000 frames per second. The reconstruction reveals the 3D distribution of RBCs and allows for cell tracking over time (**Methods, Supplementary Figure 8**). **Figure 2a** shows two separate recordings from the dorsal and ventral view, each visualizing RBCs at representative time points (red) and the vasculature network by maximum intensity projection (MIP) throughout all frames (cyan). The flowing velocity is pulsatile temporally and varies spatially in the aorta and vein (**Supplementary Videos 1, 2**). The tracking reveals the velocity distribution in 3D and highlights vessels with a high-speed flow of up to 6 mm/s (**Fig. 2a**). SLIM’s kilohertz imaging seizes the transient motion at a millisecond time scale (**Fig. 2b**), effectively eliminating the motion blur and enabling robust cell tracking compared to a lower imaging rate (**Fig. 2c**).

**Fig. 2.**
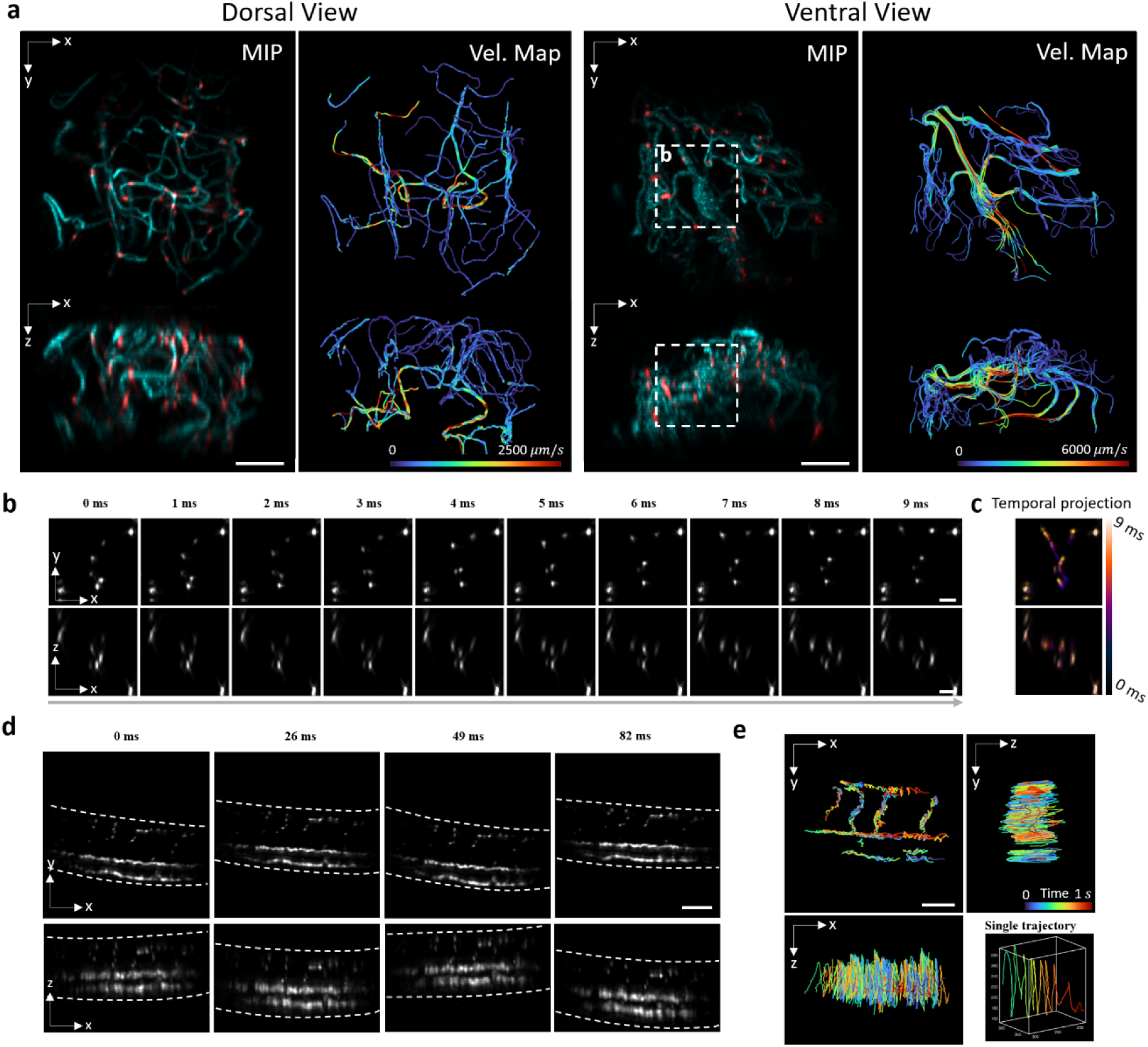
3D imaging of hemodynamics in the embryonic zebrafish brain and tail at 1,000 vps. **a.** MIPs of flowing blood cells at representative time points (red) and vascular network obtained by combining frames over time (cyan). The velocity maps show overlaid RBC trajectories color-coded by their instantaneous velocity. Two views, the dorsal view, and ventral view, are two datasets taken with differently-orientated embryos. Scale bar, 100 μm. **b.** Zoom-in time-lapse of the region labeled by the white dotted box in **a**. Scale bar, 30 μm. **c.** Temporal projection color-coded by the time visualizes the motion blur with a lower imaging speed. **d.** MIPs of a free-moving fish tail. **e.** RBC trajectories from **d** are color-coded by time. The coordinate system has been rotated so that the x-y plane shows the RBC movement perpendicular to the tail swing direction. The single trajectory on the bottom right exhibits the compound motion of a single RBC during fish swimming. Scale bar, 100 μm.

We further demonstrated the speed advantage by imaging the free-moving tail of a zebrafish without sedation. The embryo was mounted on a cover glass with its head restrained using agarose while allowing the tail to move freely in the water. SLIM captured the high-frequency tail swings without any motion blur (**Fig. 2d, Supplementary Videos 3**), maintaining its capability to track individual RBCs and revealing the compound movement that combines oscillation vertical to the tail plane and normal progression along the vessels (**Fig. 2e**). By combining with closed-loop tracking and a translational stage^16^, SLIM’s high-speed volumetric imaging holds promise for studying hemodynamics under natural conditions during locomotor behavior.

### Optical recording of membrane action potentials in medicinal leech ganglia

The development of voltage imaging has enabled neuroscientists to examine neural dynamics with a high spatio-temporal resolution. However, it has long been a challenge to capture voltage signals *in vivo* across a large volume due to the extremely fast transients and low signal-to-noise ratio. With its millisecond temporal resolution, SLIM can precisely detect spike timings across a large 3D neural network, opening avenues for mapping the intricate interaction of neuronal components and elucidating the mechanisms underlying sensory processing and behavioral generation.

As a demonstration, we loaded a voltage-sensitive dye (FluoVolt, F10488, Thermo Fisher Scientific) to a dissected ganglion from a medicinal leech.^43^ Using SLIM, we recorded the fluorescent signals with 29 sub-aperture images at 800 Hz under the illumination of an ultra-low-noise LED. Concurrently, we introduced an intracellular microelectrode for simultaneous electrophysiological stimulation and recording (**Fig. 3a**). Timing and waveforms (**Fig. 3b**) of neuronal action potentials are adequately sampled from the reconstructed 3D image sequence (**Fig. 3c, Supplementary Videos 4**). We further processed the data by correcting motion drift, manually choosing an area of interest on each cell, and averaging pixels from the corresponding cell membrane (**Methods, Supplementary Figure 9,10**). The resultant time-lapse fluorescence intensities at selected neurons are shown in **Fig.3d**. SLIM measurements agreed with the electrophysiological record in quantitative detail, including the reduction of spike amplitude when strong depolarizing currents were applied.

**Fig. 3.**
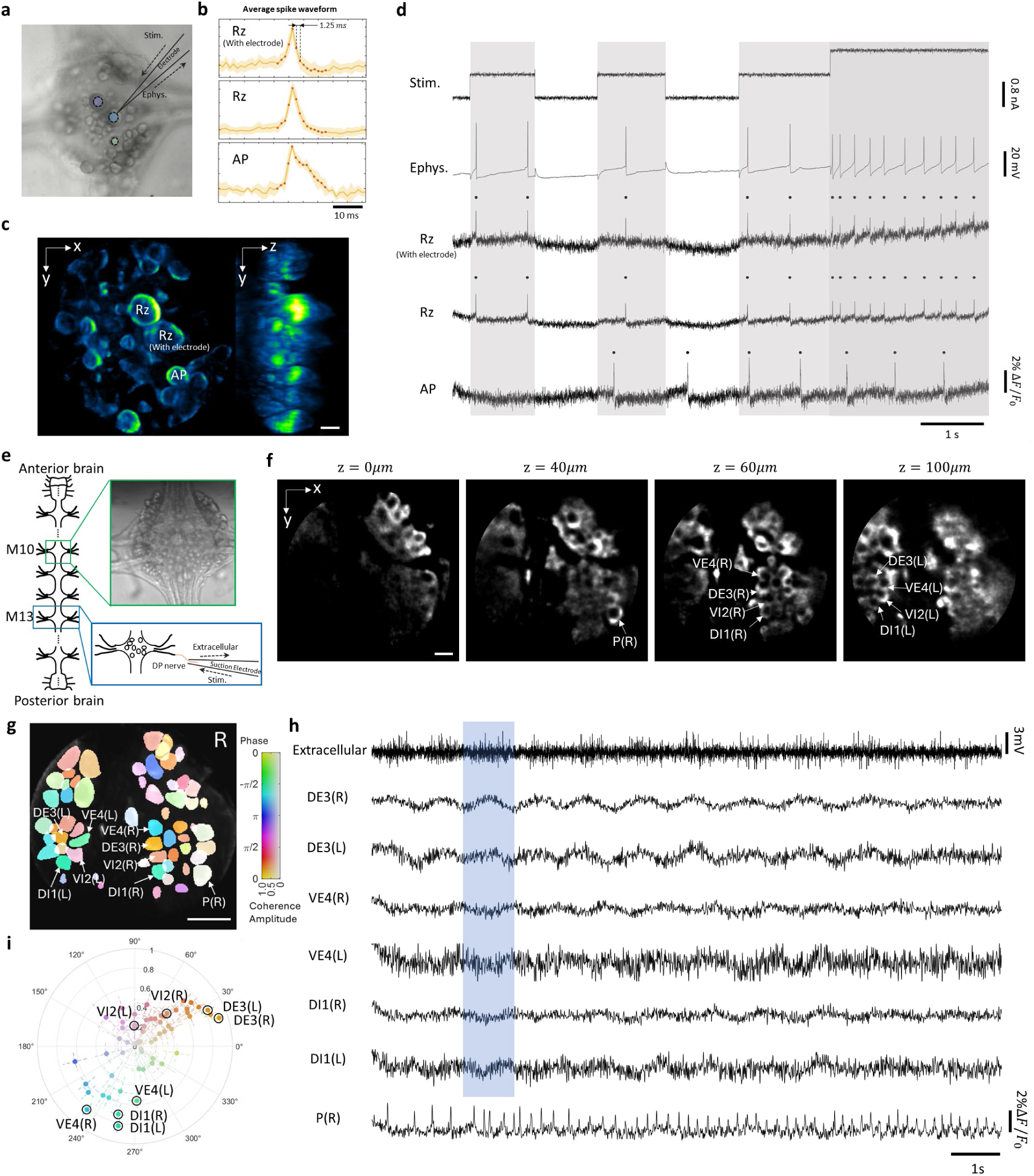
3D imaging of membrane action potentials and fictive swimming oscillation in medicinal leech ganglia at 800 vps. **a**. Brightfield snapshot of leech ganglion. Microelectrode is denoted by a dotted line, which allows for simultaneous stimulation and electrophysiological recording. **b**. Average spike waveform. The yellow area marks the standard deviation of signals. Orange dots represent temporally sampling points, with an interval of 1.25 ms. **c**. MIP of SLIM reconstruction for voltage dye fluorescent signals. Scale bar, 50 μm. **d**. Recording of stimulation current, electrophysiological readout, and optical measurements of ganglion cells: the impaled cell (top), its contralateral partner that is electrically coupled to it (middle), and an unconnected cell (bottom). Rz indicates Retzius cell. AP indicates Anterior Pagoda cell. Grey boxes represent the time window when stimulation is injected. Deeper color stands for larger stimulation. Spikes are detected and marked as black dots above the traces. **e.** A schematic of the isolated leech nerve cord, consisting of the anterior brain, midbody ganglia (circles), and the posterior brain. The voltage-sensitive dye components were applied to a midbody ganglion (M10). The brightfield reference image of selected ganglion is shown in the right. **f**. x-y cross section of SLIM reconstruction for voltage dye fluorescent signals of fictive swimming behavior. Scale bar, 50 μm. **g**. Coherence of the optically recorded signals from all cells on the dorsal surface of the ganglion with the swim rhythm. Cells used in (h) are marked. Scale bar, 120 μm. **h**. Selected electrophysiological and voltage-sensitive dye (VSD) traces of motor neurons during fictive swimming are presented. An extracellular recording from a nerve root in a posterior segment showed rhythmic bursts from dorsal motor neurons, characteristic of swimming (top). VSD signals from the dorsal surface captured the activity of dorsal and ventral inhibitory and excitatory motor neurons, specifically DI-1, DE-3, and VE-4. Blue box visualizes the synchrony of the subthreshold oscillations among neurons. **i**. The polar plot illustrates the coherence between each optical recording and the extracellular recording in the frequency range of 0.8 Hz to 1.1 Hz. The distance from the center represents the coherence magnitude, while the angle indicates the coherence phase. Error bars show confidence intervals, calculated using the multi-taper estimate method.

In a separate experiment, we used a train of electrical pulses to simulate a dorsal posterior (DP) nerve root of midbody ganglion 13 (M13), which mimics a touch to the body wall in an intact leech to elicit fictive swimming^44,45^. Using the SLIM system, we imaged the selected midbody ganglion 10 (M10) (**Fig. 3e, f**) from the dorsal side at an 800 Hz volume rate under the same illumination conditions as the previous demonstration. The nerve signal was simultaneously recorded through the suction microelectrode, which showed rhythmic dorsal motor neuron bursts characteristic of swimming (**Fig. 3h**). After manually selecting and averaging 3D region of interest for each cell, the optical fluorescence signals at selected motor neurons (dorsal and ventral inhibitory and excitatory motor neurons DI-1, DE-3, and VE-4.) and pressure sensitive cell (P2) are shown in **Fig. 3h**. The rhythmic activity characteristic of swimming (1Hz to 1.5Hz) was clearly observed in all motor neurons, consistent with previous work^43^. To characterize how cells participated in generating the swim rhythm, we calculated the magnitude and phase of coherence (see **Methods**) for each cell in the swimming oscillation band (**Fig. 3g, i**) with respect to the extracellular recording. SLIM measurement matches well with oscillatory behavior of neurons, including the overall coherence phase distribution of all cells in dorsal side and four pairs of specific motor neurons, DI-1, DE-3, VI-2 and VE-4, are very regular in their location and indeed overlapped in the measured and predicted phase maps.

### Voltage imaging from the hippocampus of behaving mice

We examined the performance of the SLIM system in awake, behaving mice. We monitored neuronal activity in the CA1 of the hippocampus through an implanted cranial window in mice expressing genetically encoded voltage indicator (GEVI) pAce^46^. Mice were imaged continuously at 800 Hz for three minutes on a treadmill set-up that used an optical rotary encoder to track movement (**Fig. 4a**). Conventional widefield microscopes suffer from a shallow depth of focus and have difficulty imaging axially distributed neuron population (**Fig. 4b**). In contrast, SLIM provides volumetric mapping of signals (**Supplementary Figure 11**, **Supplementary Table 4**), allowing for simultaneous optical measurement of neurons at different depths (**Extended Data Fig. 1**). After image reconstruction and motion correction, we extracted membrane-potential traces from multiple neuronal sources exhibiting strong speed-related action potential modulation across the imaged volume (**Fig. 4c**). We calculated the relative fluorescence change over the three-minute recording, both in signal-to-noise ratio (SNR) (**Fig. 4d, Supplementary figure 12)** and in Δ*F*/*F*_0_ (**Fig. 4e**), where SLIM’s millisecond temporal resolution provided sufficient sampling on the rising and falling slopes of transient spikes (**Supplementary figure 13**). Due to photobleaching (**Supplementary figure 14**), the amplitude of the spike waveform exhibited a gradual decay, but the SNR maintained around five across the entire recording (**Fig. 4e**). SLIM also detected subtle subthreshold membrane potential oscillations (**Fig. 4d inset**, **Extended Data Fig 2**). The observed signals predominantly show significant frequency components in the 4-10 Hz band, likely originating from theta oscillations commonly found in the hippocampus^47–49^ (**Fig. 4f, Extended Data Fig 2**). The examination over the inactive neurons and background reveals the absence of spikes and subthreshold oscillations, further confirming the fidelity of observed signals. By correlating the time-dependent firing rate of each neuron with locomotion speed, we found that the majority of neurons were positively modulated by locomotion speed (**Fig. 4g, Supplementary Figure 15s**), consistent with the previous finding^50^. Additionally, SLIM’s compressed measurement offers a unique advantage over alternative methods, providing highly efficient data bandwidth (**Fig. 4h**) and making it more accessible for long-term 3D voltage imaging across large volumes. Overall, SLIM enables 3D voltage imaging in neuron populations distributed across large volumes, with the potential to elucidate network dynamics and interactions between different cell types across layers.

**Fig. 4.**
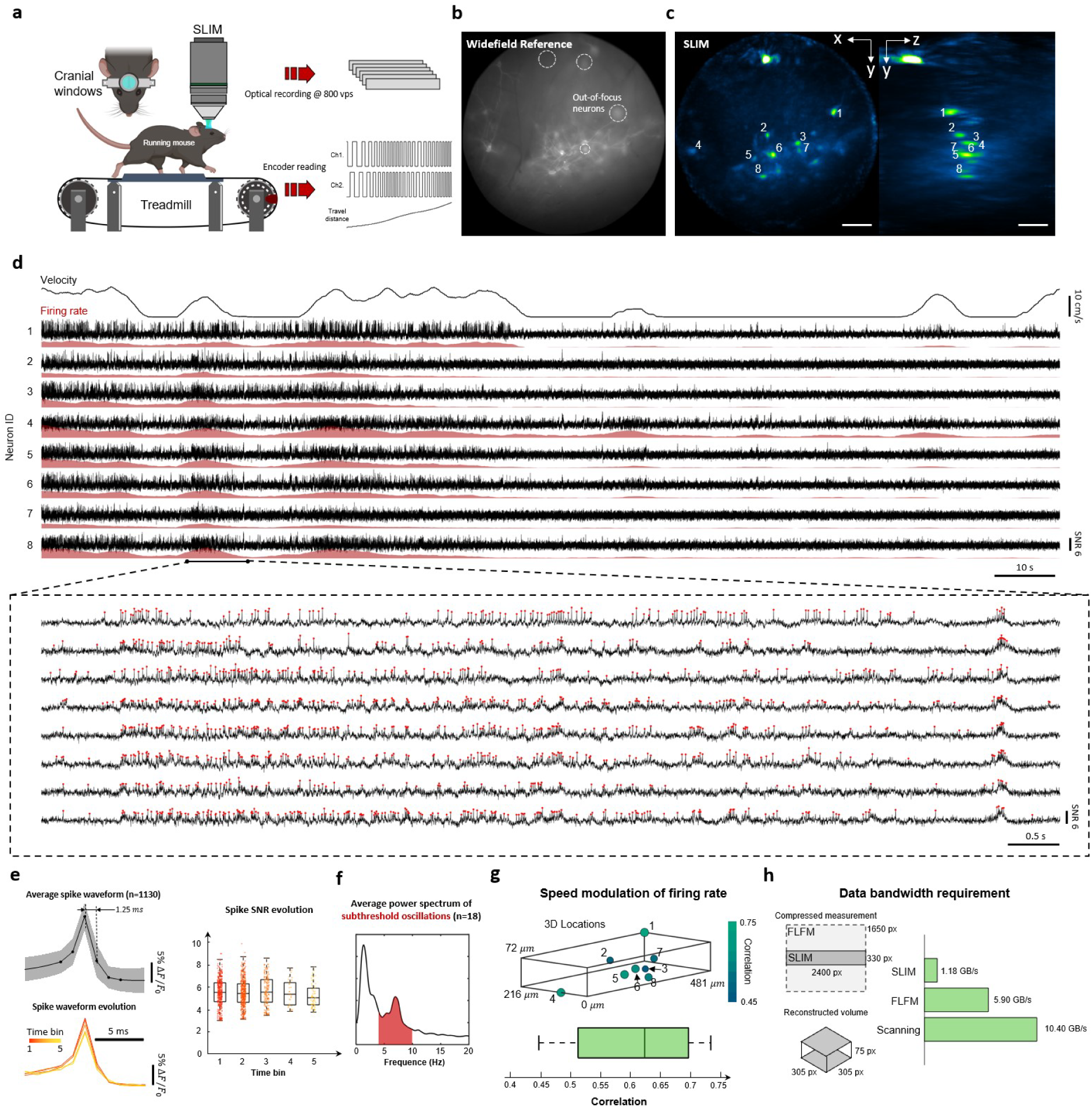
3D voltage imaging of hippocampus in behaving mice at 800 vps. **a.** Schematics of the experiment setup. The animal expressing genetically encoded voltage indicator is placed on a customized treadmill. SLIM captures volumetric fluorescent signals through cranial windows, while an optical rotary encoder on the treadmill simultaneously records the belt’s motion. **b.** Widefield reference camera captures a high-resolution, long-exposure 2D image of the targeted field of view (FOV). Dotted circles highlight out-of-focus neurons. **c.** 3D MIP of SLIM reconstruction. Scale bar, 100 μm. **d.** Detrended fluorescent signal traces over a 170-second recording, extracted from neurons labeled in panel b. The top row shows the animal’s locomotion velocity. The red-shaded curve represents neuronal firing rates. The inset zooms in on the signals within the 24–34 second window. Red dots represent detected spikes. **e.** Average spike waveform calculated from 1,130 detected spikes in neuron 1. By further dividing the signal trace into five time bins (35 seconds each), the evolution of waveform can be visualized. The spike SNR, measured as the spike magnitude over the signal’s standard deviation within each time bin, shows a gradual decrease over time. Dots represent individual spike measurements. **f.** Average power spectrum of neural signal traces (n=18), showing strong subthreshold oscillations between 4-10 Hz. See **Extended Data Fig. 2** for signal traces and their individual power spectra. **g.** Pearson correlation between firing rate and locomotion velocity reveals speed modulation of neural activity. Upper panel: 3D locations of selected neurons with correlation values encoded by color. Lower panel: distribution of correlation coefficients. **h.** SLIM’s compressed measurement significantly reduces the data bandwidth required for camera readout and storage. Raw data size for SLIM, Fourier LFM and scanning-based methods is roughly estimated based on the digital image dimensions (left panel) in 16 bits.

### Imaging of a beating embryonic zebrafish heart with scanning multi-sheet illumination

Although LFM techniques, including SLIM, offer the ability to numerically refocus to specific depths, they typically lack intrinsic optical sectioning capability. Its application is potentially hindered by the spatial resolution and reconstruction artifacts, and it favors objects with high sparseness.^51^ Here, we demonstrated that SLIM can be combined with scanning multi-sheet illumination. The synergy enables high-contrast 3D imaging of densely-labeled fluorescent objects.

We constructed a dual-light-sheet illumination module and scanned the beams using a galvo-mirror driven by a sawtooth function. Rather than synchronizing the camera exposure with the entire scan range as in previous experiments, we operated the camera at a higher rate, allowing each frame to capture a subset of depth layers of the fluorescent object (**Fig. 5a**). This approach significantly suppresses out-of-focus light and improves axial resolution in the reconstruction, as shown on fluorescent beads and zebrafish vasculature networks (*Tg*(flk:mCherry)) (**Fig. 5b,c**). On the other hand, SLIM offers an ultra-high framerate and supports simultaneous multi-plane detection. These features enable SLIM to maintain a high volume rate even within this scanning scheme.

**Fig. 5.**
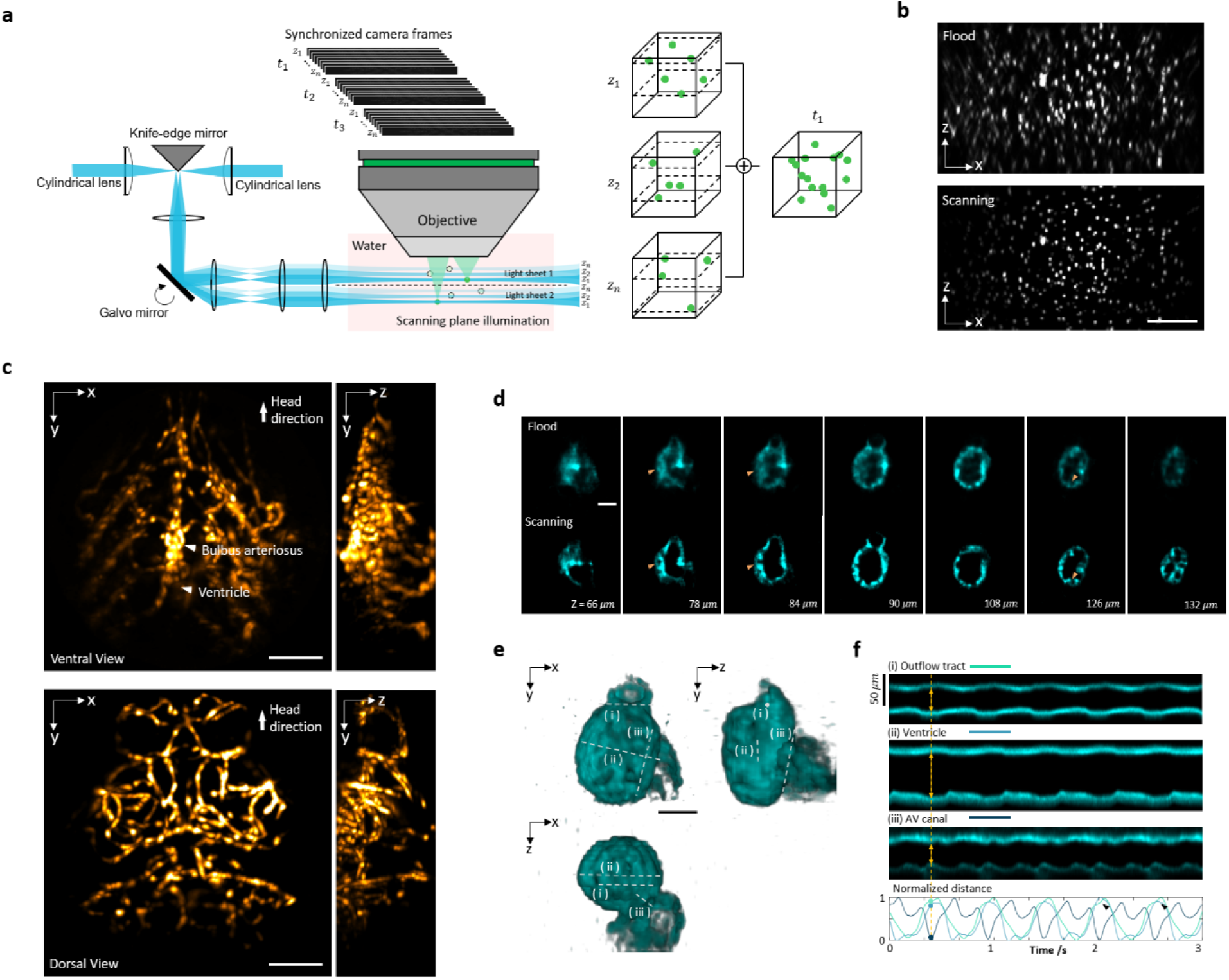
3D imaging of a beating zebrafish heart with multi-plane scanning light sheet illumination at 300 vps. **a.** Dual scanning light sheet replaces the flood illumination (*i.e.*, illuminating entire sample volume). In synchrony with the light sheet, the camera captures multiple frames at different scanning positions, each reconstructing two layers of the entire volume. By combining all measurements in one scan cycle, a 3D volume is synthesized for that time point. **b.** The x-z MIPs of fluorescent beads show the enhancement of image contrast and axial resolution. Scale bar, 100 μm. **c.** The structural images of the vasculature network in an embryonic zebrafish. Scale bar, 100 μm. **d**. Comparison of x-y cross-section images between scanning light sheet illumination and flood illumination on cardiomyocytes in the zebrafish heart. The orange arrows mark the muscle structures that are clearly resolved in scanning mode but challenging to flood illumination. Scale bar, 50 μm. **e.** The 3D rendering of myocardium at a representative time point. Scale bar, 50 μm. **f.** Kymographs calculated by sampling the time-dependent distance of the cavity along the white dotted line in **e**. The black arrows indicate the beat-to-beat variance of cardiac contraction. Scale bar, 50 μm.

We demonstrated this acquisition scheme by imaging a beating zebrafish heart (*Tg*(cmlc2:GFP)) at 300 vps (**Fig. 5d, Supplementary Videos 5**). This is achieved by scanning the dual light sheets at 300 Hz, synchronized with camera recording at 4,800 fps (8-bit speed mode, see **Methods**). This setup allowed us to reconstruct the heart with 30 planes across 200 μm depth range with the microstructures like ventricular trabeculation clearly delineated (**Fig. 5d,e**). The enhanced spatial resolution and contrast offer the potential for accurate segmentation of the heart chamber’s geometry, facilitating cardiac studies, such as regional myocardial contractility analysis^40^ and computational fluid dynamics (CFD) for hemodynamic forces simulation^52^. While current LFM cardiac imaging is mostly demonstrated on sparse markers like cardiomyocyte nuclei^19,20,53^ and blood cells^20,21,54^, SLIM with scanning multi-sheet illumination proves effective in resolving the densely labeled muscle tissue. It provides high 3D imaging speed to capture the beating heart in real time and outlines the time-dependent chamber dimension to detect beat-to-beat variations (**Fig. 5f**).

## Discussion

We presented SLIM as an innovative snapshot 3D detection method that addresses the pressing need for high-speed volumetric microscopy operating at kilohertz speeds. SLIM accomplishes this by capturing a condensed representation of the original light field using a compact ROI on the sensor. The sampling strategy is grounded in the principle that the inherent spatio-angular correlation in the light field can be exploited to recover signals from compressive measurement^24,25,27–29,31,55^. As a compressed sensing method, successful reconstruction with SLIM assumes sample sparsity (**Supplementary Note 2**). However, SLIM has been successfully demonstrated across a range of applications, including hemodynamics, neural imaging, cardiovascular imaging, and bacterial dynamics (**Supplementary Figure 16**), showcasing its versatility and robustness.

SLIM’s kilohertz 3D imaging speed, rarely provided by existing methods and often entailing significant design tradeoffs and demanding hardware requirements^9,23^, presents new opportunities to investigate millisecond-scale dynamics in emerging fields like voltage imaging. It is universally adaptable to the vast majority of CMOS sensors, which generally allow higher frame rates at reduced readout pixel rows. While we set kilohertz as a milestone for 3D fluorescence microscopy, SLIM has the potential to achieve tens and even hundreds of thousands of vps with current ultra-fast cameras^56^.

Quantum efficiency and readout noise are critical parameters that determine a sensor’s sensitivity in low-light imaging. Unfortunately, these factors degrade significantly in the ultra-fast cameras currently available (**Supplementary Table 1**). Another important factor, the full well capacity, is also frequently sacrificed together with bitdepth in trade for framerate. It becomes a relevant limitation when the fluorescence change (ΔF/F₀) is extremely low (*e.g.* our leech ganglion imaging) and a high pixel brightness is required to provide an acceptable SNR^7^. Moreover, the extended-time imaging in animal behavior study often demands a significant number of frames to be recorded, which becomes impractical for ultra-fast cameras that rely on limited on-board storage (**Supplementary Table 1**). Although this could be potentially solved by a camera array with a dedicated computer cluster for data handling^57^, there are considerable technical and financial challenges in integrating high performance sensors into an array. In contrast, SLIM presents an accessible solution to transform a single commonly used sCMOS into a kilohertz 3D imaging tool, as demonstrated in various demanding applications, including voltage imaging in the mouse brain. SLIM offers a snapshot acquisition that effectively addresses the trade-off between pixel exposure time and volumetric frame rate encountered in conventional scanning-based 3D optical microscopy techniques. This unique approach gives SLIM distinct advantages in terms of photon efficiency and signal-to-noise ratio (SNR), especially beneficial for high-speed imaging of weak fluorescence.

SLIM compresses images only along the vertical axis and redistributes the information to the horizontal axis, which retains full sampling power through image rotation. As a result, SLIM can reconstruct the FOV of a FLFM that occupies the full sensor area, achieving comparable spatial resolution, provided the sample’s sparsity allows. This design is specifically tailored to utilize a low-format rectangular sensor, marking a fundamental departure from existing compressive light field imaging^24–30,55^. The latter retrieves light fields at the same or lower resolutions than the multiplexed measurement and suffers from a linear reduction in FOV and pixels as it crops the sensor ROI. Moreover, SLIM does not require multiple shots^24^ or learning on a sparse basis prior to reconstruction^26^. It also shows excellent scalability in various ROI sizes. As demonstrated in the numerical simulation (**Supplementary Note 2**), we can reduce the vertical scaling ratio to 0.1 or even 0.04, and still reconstruct the same FOV with as few as 160 or 64 vertical pixels in the raw measurement. As long as the sample sparseness permits, it can tolerate this extreme narrow sensor size and further increase its space-bandwidth-framerate product by using a faster recording speed. SLIM can transform an existing FLFM^16,35,51,58^ into a high-speed 3D imager with a significantly higher frame rate. Its performance approximates FLFM when the targeted dynamics exhibit sufficient spatiotemporal sparseness (**Supplementary Note 2**). However, as with all compressive detection systems, degradation is expected when handling dense signals with complex structures (**Supplementary Note 2**). Moreover, like FLFM, SLIM faces challenges such as the missing cone problem, compromised spatial resolution, depth-variant performance, and limited considerations for tissue scattering and lens aberrations. In addition, our current reconstruction assumes no occlusions in the scene^27,29^. These factors constrain its applicability in complicated intravital environments. We have shown that multi-sheet scanning offers a solution by trading imaging speed for improved optical sectioning, extending SLIM’s applicability to densely-labeled tissue imaging. The refocusing capability within a largely extended depth of field makes SLIM compatible with various 3D illumination structures. Additionally, the literature presents several strategies to enhance SLIM’s performance, such as background rejection by hardware^18^ and computation^59^, multi-focus optics for extended DoF^16,60^, and sparsity-based resolution enhancement^51,55^. Furthermore, ongoing advancements in data-driven reconstruction algorithms, particularly physics-embedded deep learning models^20,21,53,61^, hold great promise for addressing the ill-posed inverse problems associated with limited space-bandwidth and compressive detection in SLIM. These developments are expected to significantly enhance SLIM’s capabilities and broaden its utility across diverse imaging scenarios.

## Acknowledgements

We would like to thank Yuan Dong and Yaran Zhang at UCLA for their assistance in zebrafish experiments. We acknowledge the David Geffen School of Medicine at UCLA for providing the Zebrafish Core Facilities. This work was supported by the following grants: National Institutes of Health (NIH) (R01HL165318, RF1NS128488, R35GM128761, R01AI102584, R01HL129727, R01HL159970, 5T32HL144449). W.C.S. was supported by NSF Graduate Research Fellowship Program (DGE-1650604, DGE-2034835) and Ruth L. Kirschstein National Research Service Award “Multidisciplinary Training in Microbial Pathogenesis” (T32AI007323).

## Author contributions

Z.W. and L.G. conceived the idea. L.G., D.A.W., R.L., G.C.W., T.K.H. and P.G. oversaw the project. Z.W. constructed the microscopes and developed the reconstruction algorithm. R.Z., D.A.W. and Z.W. prepared medicinal leech samples and electrophysiology study. D.E. and L.S. performed mouse surgery. Z.W., R.Z., A.P., and J.W. bred the zebrafish. C.K.L. and W.C.S. prepared bacteria samples. W.K. fabricated the lenslet array. Z.W., R.Z., D.A.W., D.E., L.S. O.B. and E.Z. collected the imaging data. Z.W. and R.Z. processed and analyzed the data. Z.W., R.Z., and L.G. wrote the manuscript. All authors reviewed, edited, and consulted on the manuscript text.

## Methods

### Hardware setup

For SLIM setup using selective volume side-illumination, the detection features a 20X water-dipping objective (N20X-PFH, Olympus XLUMPLFLN 20X, 1.0 NA). A 4F relay system (AC508-180-A, AC508-200-A, Thorlabs) forms a conjugate plane of the objective’s back pupil, accommodating a customized dove prism and a spherical lenslet array. The dove prism (aperture length: 1.3 mm, material: H-K9L, fabricated by Changchun Sunday Optics) is positioned anteriorly to the plano-convex lenslet (aperture diameter: 1.3 mm, focal length: 36 mm, material: PMMA, fabricated in-house). Each pair generates a rotated sub-aperture image with a magnification of 3.6X and NA of 0.065. In total, 29 pairs are utilized and securely housed in 3D-printed mechanical holders (refer to **Supplementary Figure 1** for detailed designs of the prism, lenslet, and holder). Our anamorphic relay system comprises a spherical achromat doublet (ACT508-250-A, Thorlabs) and two orthogonally oriented cylindrical achromat doublets (ACY254-250-A, ACY254-50-A, Thorlabs). The back focal planes of two cylindrical lenses are co-located, producing an image with an anisotropic scaling factor (**Supplementary Figure 2**). A sCMOS camera (Kinetix, Teledyne) captures the final image, with a 320 × 3200 pixels ROI covering all 29 sub-aperture images, or a 200 × 3200 pixels ROI for 19 sub-aperture images. The maximal readout speeds for two ROIs are 830 fps and 1,326 fps in 16-bit dynamic range mode and 4,790 fps and 7,476 fps in 8-bit speed mode. The 29 sub-aperture configuration collects around 50% more light than the 19 sub-aperture one.

The illumination sources include blue and green continuous lasers (MBL-FN-473-500mW and MGL-III-532-300mW, CNI Laser) and an ultra-low-noise blue LED (UHP-T-470SR, Prizmatix). For scanning light sheet setup, we use a knife-edge mirror (MRAK25-G01, Thorlabs) to combine two beams with adjustable spacing and a galvo-mirror (GVS011, Thorlabs) to scan them together. Planar illumination is formed perpendicular to the detection axis by a cylindrical lens and a dry objective (RMS4X-PF, Olympus 4X, 0.13 NA). We use a sawtooth function to drive the galvo-mirror. In synthetic volume illumination configuration, we block one light sheet beam. The camera is then triggered at the beginning of the sawtooth waveform and exposed for the entire scan. In scanning plane illumination configuration, we use two light sheet beams and trigger the camera multiple times during a scan. The static LED setup shares the same illumination objective and perpendicular geometry. We build a Koehler illumination system and use an adjustable slit as a field aperture. The conjugate image of the slit is relayed to the sample, and the slit controls the depth range of the beam. The LED provides ultra-stable illumination power and thus suppresses excitation source noise during our voltage imaging experiments.

For SLIM setup using widefield epi-illumination, we use a 16X water-dipping objective (16X Nikon CFI LWD Plan Fluorite Objective, 0.80 NA). The back pupil relay system uses a pair of doublets (AC508-150-A, Thorlabs). The dove prism, lenslet array, and anamorphic relay system are identical to the selective volume side-illumination setup. An ultra-low-noise blue LED (UHP-T-470SR, Prizmatix) in a Koehler configuration is used for epi-illumination.

See **Supplementary Figure 4, 5** for system schematics, **Supplementary Table 3** for components list, and **Supplementary Table 4** for configurations used in different experiments.

### Image formation and reconstruction

The light field of the fluorescent sample is acquired by dividing the objective’s back pupil with a lenslet array and recording a group of sub-aperture images. Depending on their sub-aperture locations, they display disparity, that is the distinct displacement shown by the same signal. After calibrating the displacement at every axial position, the formation of each sub-aperture image can be modeled as a sum of laterally shifted depth slices. We replace the shifting operator with a convolution with PSF to account for both the diffraction and displacement. A dove prism is a truncated right-angle prism and used to rotate the incident beam. The rotation of the prism around its longitudinal axis causes the beam to rotate at twice the rate of the prism’s rotation. By placing a dove prism array in the infinity space between the objective and lenslet array, we apply varying in-plane rotations to sub-aperture images. Finally, we adopt an anamorphic relay system to introduce anisotropic scaling to the image array: we de-magnify (squeeze) the image in the direction perpendicular to the camera read-out axis while maintaining the original scale in the other direction (**Supplementary Figure 2**). This one-axis scaling and the aforementioned in-plane rotation are both directly applied to the 3D fluorescent image in our model.

Given a 3D fluorescence distribution *O*(*x*, *y*, *z*) and system PSF, the formation of camera measurement *I*(*x*, *y*, *v*) can be modeled as:

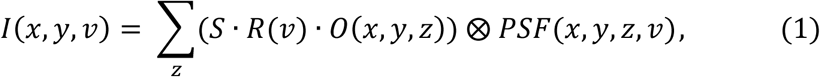

where *v* is the index of sub-aperture, ⊗ represents the 2D convolution, *R*(*v*) applies sub-aperture-dependent rotation from the dove prism, and *S* introduces image scaling from the anamorphic relay system.

The volume reconstruction algorithm was derived from Richardson-Lucy deconvolution^15,16,35,62^ (**Supplementary Note 3**). Based on the forward model (1), the 3D fluorescence distribution *O*(*x*, *y*, *z*) is iteratively solved from the camera measurements *I*(*x*, *y*, *v*) and empirical point spread functions *PSF*. The rotation angles and image scaling ratio are pre-calibrated as known priors in the reconstruction. The measurement patch *I*(*x*, *y*, *v*) is cropped from the raw sensor image according to the center location of each sub-aperture image. We experimentally measure the point spread functions by imaging a sub-diffraction fluorescent bead. The point source is placed in the middle of FOV and axially scanned over a broad 600 μm depth range with a 2 μm step size using a motorized translation stage. The actual axial range and step size in the reconstruction depends on the specific experiments. The PSF is assumed to be spatially invariant within each sub-aperture image.

With our implementation, each measurement patch *I*(*x*, *y*, *v*) has a resolution of 301 × 61 pixels. The numbers of channels (*v*) and axial slices (*z*) are configured based on the targeted framerate and depth range/step size. For example, with 19 sub-aperture measurements to reconstruct a volume of 305 × 305 × 151 pixels, the deconvolution takes around 30 seconds via eight iterations using a desktop computer with a modest GPU (Nvidia RTX 3070).

### Fluorescent beads imaging and resolution characterization

Fluorescent microspheres of 1 μm (F13082, ThermoFisher) were embedded in 1% low-melting agarose and injected into a transparent fluorinated ethylene propylene tube for imaging. To acquire experimental PSFs, the solution was highly diluted to keep only one bead in the FOV. For spatial resolution quantification, the beads were randomly distributed in space, and we measured the full width at half maximum (FWHM) of the bead image at various locations in the FOV (**Supplementary Figure 3**). We also performed two-point resolving experiments by reconstructing synthetic measurements that added two sequentially captured frames. We translated the bead laterally and axially at varying distances in these two frames. The resolution was then indicated by the resolvable set with the least distance (**Supplementary Figure 3**).

### Fish husbandry and imaging

Transgenic zebrafish lines *Tg(gata1a:dsRed)*, *Tg(flk:mCherry)*, and *Tg(cmlc:GFP)* were used in our experiments for imaging blood cells, endothelial cells, and myocardium, respectively. Embryonic fish were maintained at three days post-fertilization in standard E3 medium, which was supplemented with extra 1-phenyl 2-thiourea (Sigma Aldrich) to inhibit melanogenesis at day one. For brain hemodynamics and cardiac imaging, the larvae were anesthetized with tricaine (3-aminobenzoic acid ethyl ester, Sigma Aldrich) and immobilized in 1% low-melting-point agarose inside a fluorinated ethylene propylene tube before imaging. For tail experiments, the larvae were first positioned on cover glass before the heads were fixed by 3% low-melting-point agarose. Immediately after the agarose solidified, the sample was immersed in a water chamber. The imaging starts after visually confirming the unconstrained movement of the tail. All the experiments were performed in compliance with and with the approval of a UCLA IACUC protocol.

### Blood cell velocimetry

3D cell tracking was performed by ImageJ and the plugin Trackmate^63,64^. We used the DoG (Difference of Gaussian) blob detector and the LAP (Linear Assignment Problem) tracker for all experiments. The spot positions and trajectory properties were exported to MATLAB for further analysis and visualization. We removed the local oscillation from the tracking result by fitting a center line of each trajectory with a smooth spline. The tangential velocity was calculated by projecting the cell displacement in adjacent frames onto the center line. The trajectories were then color-coded by the tangential velocity to display the blood flow speed along the vessels. The tracking accuracy (**Supplementary Figure 8**)

### Leech sample preparation

Medicinal leeches (*Hirudo verbana*) obtained from leech.com were housed in artificial pond water maintained at 15°C. Detailed dissection procedures have been described before.^43,65^ Briefly, an adult leech was anesthetized in ice-cold leech saline and an individual segmental ganglion (M10 or M11) was dissected out. The ganglion was pinned down ventral side up on a rectangular-shaped flat substrate made of Polydimethylsiloxane (PDMS) (Sylgard 184, Dow Corning). After removing the sheath that covers the ganglion, a voltage-sensitive dye^66^ (FluoVolt, ThermoFisher) was bath-loaded using a peristaltic pump. The sample was placed under the detection objective of the SLIM system for imaging. In swim experiments, the entire nervous system was dissected out, except for the cephalic ganglia. Segmental ganglion M10 or M11 was desheathed as before. Additionally, the dorsal posterior nerves (DP1) of ganglion M13 or M14 were exposed for extracellular stimulation and recording with a suction electrode. Nerve stimulation in these caudal ganglia is a well-established method for eliciting fictive swimming.

### Leech ganglion electrophysiology and imaging

Glass microelectrodes (20–50 MΩ) were filled with a recording solution of 3 M potassium acetate and 60 mM potassium chloride. After penetrating the membrane of a cell of interest, small negative holding currents were injected to ensure stability. Intracellular electrophysiology used Neuroprobe amplifiers (Model 1600; A-M systems). Membrane voltage and electrode current were digitized along with the camera trigger signal at 10 kHz using a 16-bit data acquisition board (NI USB-6002; National Instruments). We used the camera triggers as time stamps to align recorded frames with electrophysiological data. Extracellular electrophysiology used a custom built differential amplifier that allowed for rapid switching between stimulation and recording.

After image reconstruction, we corrected for sample movement by running a 2D registration and demotion between adjacent frames using a modified version of SWiFT-IR^67^. The 3D ROIs were then manually defined for each neuron. The optical readout *F*_*t*_was calculated by averaging the pixel intensities in the ROI and normalized by the temporal baseline: *F* = (*F*_*t*_ − *F*_0_)/*F*_0_, where *F*_0_ is the temporal mean value. To detect spikes from the optical signal, we detrended the trace *F* by subtracting its median-filtered version (window size, 50 ms). It was then binarized by a Schmitt trigger, and a peak detection was performed to locate the voltage spikes.

### Coherence analysis method

We employed multitaper estimation techniques^68^ to calculate the power spectral densities denoted as *S*_*opt*_(*f*) for the optical signals and *S*_*ref*_(*f*) for the reference signal, and their coherence, denoted as *C*(*f*). The use of multiple tapers ensures a more balanced weighting across all regions of a record, in contrast to the central bias introduced by a single taper. Additionally, this approach allows for the estimation of the standard deviation of the spectral estimates within a single trial. The spectral measures are defined by 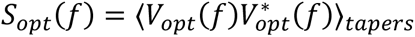. The brackets denote an average over all trials and tapers, specifically, 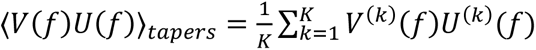, where K is the number of tapers.*V*^(*k*)^(*f*) is the discrete Fourier transform of the product *w*^(*k*)^(*t*)*V*(*t*), which refers to the time-domain optical signal *V*(*t*) multiplied by the kth taper *w*^(*k*)^(*t*). The discrete prolate spheroidal sequences (Slepian sequences) are used as tapers to minimize power leakage between frequency bands. In fictive swimming datasets (T = 12s), we selected extracellular recording as reference signal and used 10 tapers (K=10) for spectral coherence analysis. Additional processing steps, including demotion and detrending, were applied prior to the coherence analysis.

Standard deviations of the coherence are reported as jackknife estimates within single trials^69^. In this method, the variance of a dataset with K independent estimates of a quantity is calculated by sequentially deleting each estimate and computing the variance over the K resulting averages. The “delete-one” averages of coherence, denoted as *C*_*j*_(*f*) where j is the index of the deleted taper, are given by 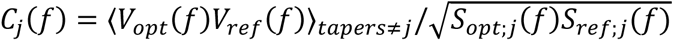, Where 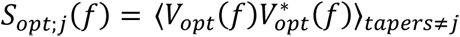

The standard deviation of the magnitude of *C*(*f*) is then computed as 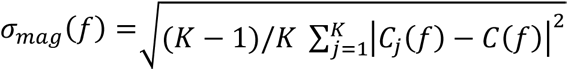. The variance estimate for the phase of is determined by comparing the relative directions of the delete-one unit vectors. The standard deviation is computed as 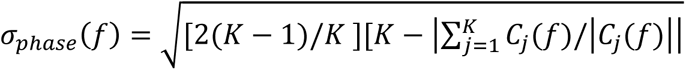.

### Behaving mouse sample preparation

All experiments were conducted according to the National Institute of Health (NIH) guidelines and with the approval of the Chancellor’s Animal Research Committee of the University of California, Los Angeles. Mice were anaesthetized with isoflurane (5% for induction, 1–2% (v/v) for maintenance). The depth of anesthesia was monitored continuously and adjusted when necessary. After induction of anesthesia, the mice were fitted into a stereotaxic frame (Kopf), with their heads secured by blunt ear bars and their noses placed into an anesthesia and ventilation system (David Kopf Instruments). Body temperature was kept at 37 °C with a feedback-controlled heating pad (Harvard Apparatus). Mice were administered 0.05 ml lidocaine (2%; Akorn) subcutaneously as a local anesthetic before surgery. The surgical incision site was cleaned three times with 10% povidone-iodine and 70% ethanol. After removing the scalp and clearing the skull of connective tissues, we drilled a hole above the virus injection location. Then, we injected GEVIs to the CA1 of the hippocampus, with coordinates ML: ±1.8mm, AP: −2mm, DV: −1.3mm from Bregma. One Chrna2-Cre+ mouse was injected with Cre-dependent GEVI pAce (AAV-DJ-CAG-DIO-pAce-kv2.1; titer, 2.6×10^12^ viral genomes per mL), allowing for imaging of neuron populations expressing nicotinic acetylcholine receptor alpha2, a specific marker for oriens lacunosum-moleculare (OLM) interneurons^70^. Another wild-type mouse was injected with a cocktail of the GEVI pAce (AAV9-EF1a-DIO-pAce-Kv-WPRE, titer, 2.1×10^13^ viral genomes per mL) and a principal-cell specific Cre promoter (pAAV1-CamKII-Cre, Addgene, 105558; titer, 1.9×10^13^ viral genomes per mL) allowing for imaging of excitatory pyramidal neurons. After the termination of viral injection, a circular craniotomy (3 mm diameter) was made around the injection site. Dura over the exposed brain surface was removed and the cortical tissue above the dorsal CA1 was carefully aspirated using a 27-gauge blunt needle. Buffered artificial cerebrospinal fluid (7.888 g NaCl, 0.372 g KCl, 1.192 g HEPES, 0.264 g CaCl2, 0.204 g MgCl2 per 1000 mL milipore water) was constantly applied throughout the aspiration to prevent desiccation of the tissue. The aspiration ceased after partial removal of the corpus callosum and bleeding terminated, at which point a 3-mm titanium ring with a glass coverslip attached to its bottom was implanted into the aspirated area and its circular flange was secured to the skull surface using vetbond (3M). A custom-made lightweight metal head holder (headbar) was attached to the skull posterior to the implant. Cyanoacrylate glue and black dental cement (Ortho-Jet, Lang Dental) were used to seal and cover the exposed skull. During recovery mice were administered carprofen (5 mg per kg of body weight) for 3 days as a systemic analgesic and amoxicillin antibiotic (0.25 mg/ml in drinking water) through the water supply for 7 days.

### Behaving mouse imaging

Mice were imaged at least three weeks after surgery on a treadmill set-up that used an optical rotary encoder to track movement for three minutes with epochs of locomotion and stationary behaviors. SLIM was configured to image at 800 Hz with a 1230 μs exposure time for each frame, under a widefield epifluorescence illumination (UHP-T-470SR Prizmatix). The camera was externally triggered. A data acquisition board (NI USB-6002; National Instruments) recorded both the camera trigger signals and the two-channel rotary encoder outputs simultaneously at 10 kHz, allowing for alignment between optical readout and animal locomotion.

The three-minute raw data from the camera (dynamic range mode, 16 bits) were continuously streamed to the host computer through a PCIe Gen 3 x16 interface and saved on four SSDs (Samsung 970 Pro 1TB) in RAID-0 configuration via a controller (HighPoint SSD7101A-1). We also tested direct streaming to a single M.2 NVMe SSD (Samsung 970 Pro 1TB). Both storage configurations enabled non-missing-frame recording at 800 Hz for up to ten minutes, thanks to SLIM’s compressed data load. However, performance may decline after repeated acquisitions without sufficient intervals for SSD cache recovery. Additionally, extended continuous recording is limited by fluorescence photobleaching (**Supplementary Figure 14**). See **Supplementary Table 4** for the illumination power in mice imaging experiments.

Each raw measurement contained 29 sub-aperture images, and we reconstructed a 3D volume with an axial range from −296 to 296 μm at a step size of 8 μm by eight iterations. The resulted image stack had a resolution of 305×305×75 pixels and took around 1 second on average after distributing the entire time sequence (∼3 minute, 144000 frames) to 24 parallel workers (Parallel Computing Toolbox, MATLAB) in our workstations (GPUs: Nvidia RTX 2080, 3070 and 3090).

Similar to leech ganglion experiments, we corrected sample movement by SWiFT-IR^67^ and manually defined 3D ROIs for each neuron candidates. The optical readout was calculated by averaging the pixels in the ROI and normalized by the temporal mean value. The spikes are detected by detrending the signals with a moving median filter (window size, 125 ms), binarizing by a Schmitt trigger, and a peak detection. The signal traces visualized in the figures (**Fig. 4, Extended Data Figure 1, 2, Supplementary Figure 15**) used larger filter window size (one second) to keep the subthreshold oscillations. We calculated the ratio between the signal and the standard deviation of entire trace to represent the SNR. However, in the analysis of spike waveform evolution, the standard deviations were calculated for each individual time bin (35 seconds), respectively. The subthreshold oscillations (**Extended Data Figure 2**) were analyzed by a band-pass filter (4-10 Hz) on the detrended signal (median filter window size, one second). To calculate the firing rate and animal locomotion velocity, we counted the number of spikes and rotary encoder pulses in the same sliding window (window size, five seconds).

#### *Vibrio cholerae* strain and preparation conditions

Wild Type Vibrio cholerae O1 El Tor strain A1552 were grown in lysogeny broth (LB) at 30°C with 200rpm shaking for 22 hours to stationary phase. Cell cultures were then diluted 1:1000 in 1mL of 2% LB media pre-mixed with 0.1ug/mL FM1-43FX membrane stain (Life Technologies F35355). 50uL of the stained cell culture was then loaded into a transparent FEP tube and sealed for imaging.

#### *Vibrio cholerae* swimming metrics

Cell trajectories were analyzed for their Radius of Gyration (RoG) and Mean Squared Displacement (MSD) slope parameters. The Radius of Gyration(RoG) quantifies the average movement of a trajectory over time which is defined as: 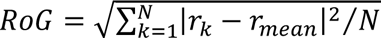, where *r*_*k*_ is the *k*^*t*ℎ^position of the cell, *r*_*mean*_ is the mean position and N is the total number of tracking point. The MSD slope is a commonly used metric in colloidal studies and biophysics to determine the mode of displacement of particles followed over time. Actively swimming cells will have an MSD slope value in the super-diffusive regime (MSD slope > 1.4) while non-swimming cells will exhibit diffusive or sub-diffusive behavior (MSD slope ≤ 1). We used Msdanalyzer^71^ for MSD slope calculation for all trajectories (https://github.com/tinevez/msdanalyzer).

### Image reconstruction with scanning multi-plane illumination

The axial positions of plane illumination during a scan were calibrated using fluorescent beads. For each scanning step, we reconstructed the bead measurement and localized two slices exhibiting the highest image contrasts. In the following imaging experiments, the same slices were sampled from the reconstruction and constituted a new stack with all other scanning steps. During high-speed scans, the sawtooth function that drives the galvo mirror could suffer from the limited bandwidth of the waveform generator. We removed the measurements at the beginning/end of the scan if repeating abnormalities (that happened at every cycle) were observed during the calibration. For image reconstruction of each scanning step, we treated it as the same reconstruction problem under volumetric illumination.

## Data availability

The data generated and analyzed in this study are available from the corresponding author upon request.

## Code availability

Codes for 3D reconstruction are available at https://github.com/aaronzq/SLIM. Codes for data analysis and post-processing used in the current study are available from the corresponding authors upon request.

---

**Extended Data Fig. 1.**
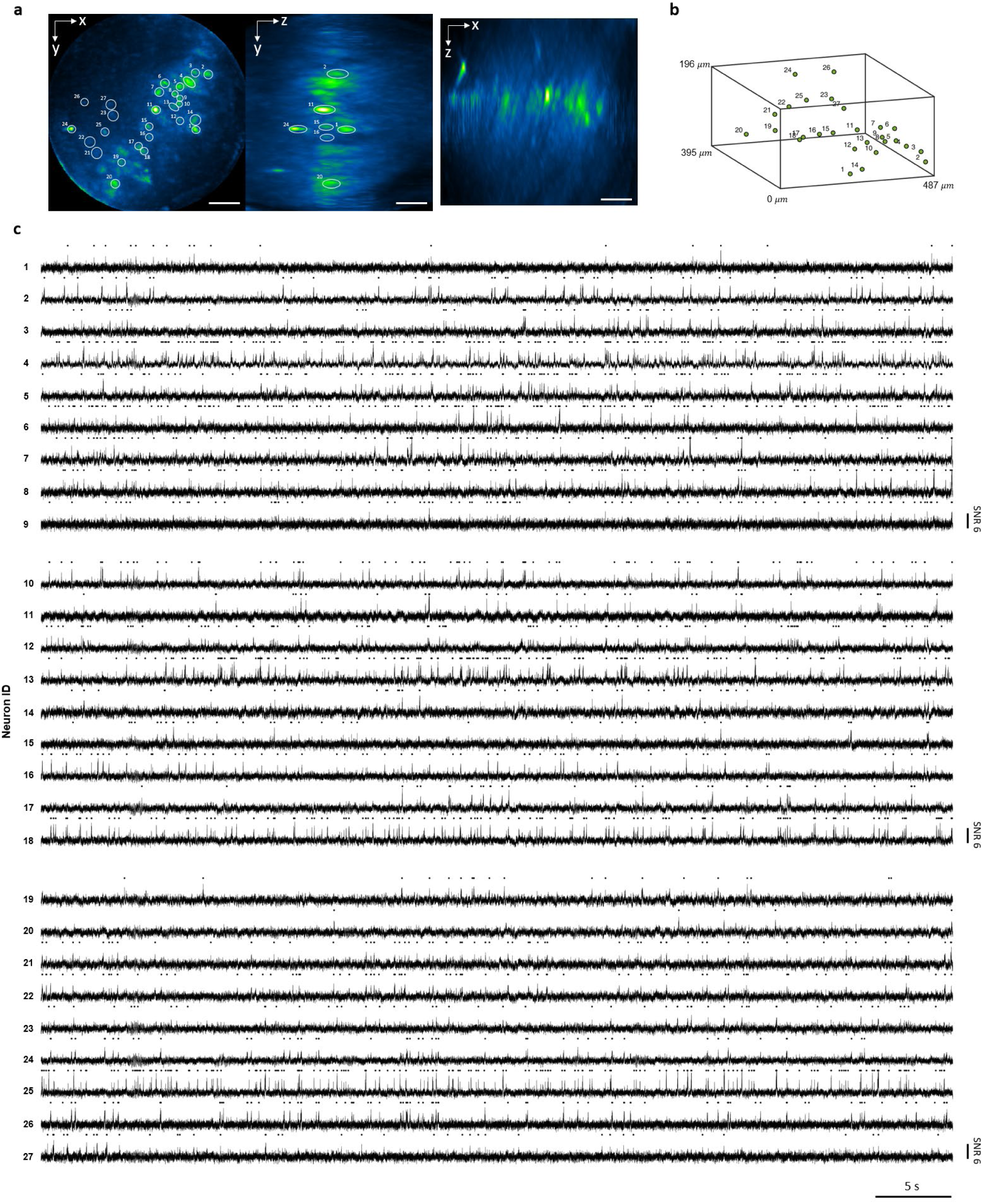
3D voltage imaging of hippocampus pyramidal neurons in awake mice. **a.** MIP of SLIM reconstruction of the 3D located neurons. Neuron indices are partially labeled in y-z view to avoid cluttered markers. Scale bar, 100 µm. **b.** 3D distribution of neuron center locations and their corresponding index. **c.** Detrended signal traces and the detected spikes labeled in black dots.

**Extended Data Fig. 2.**
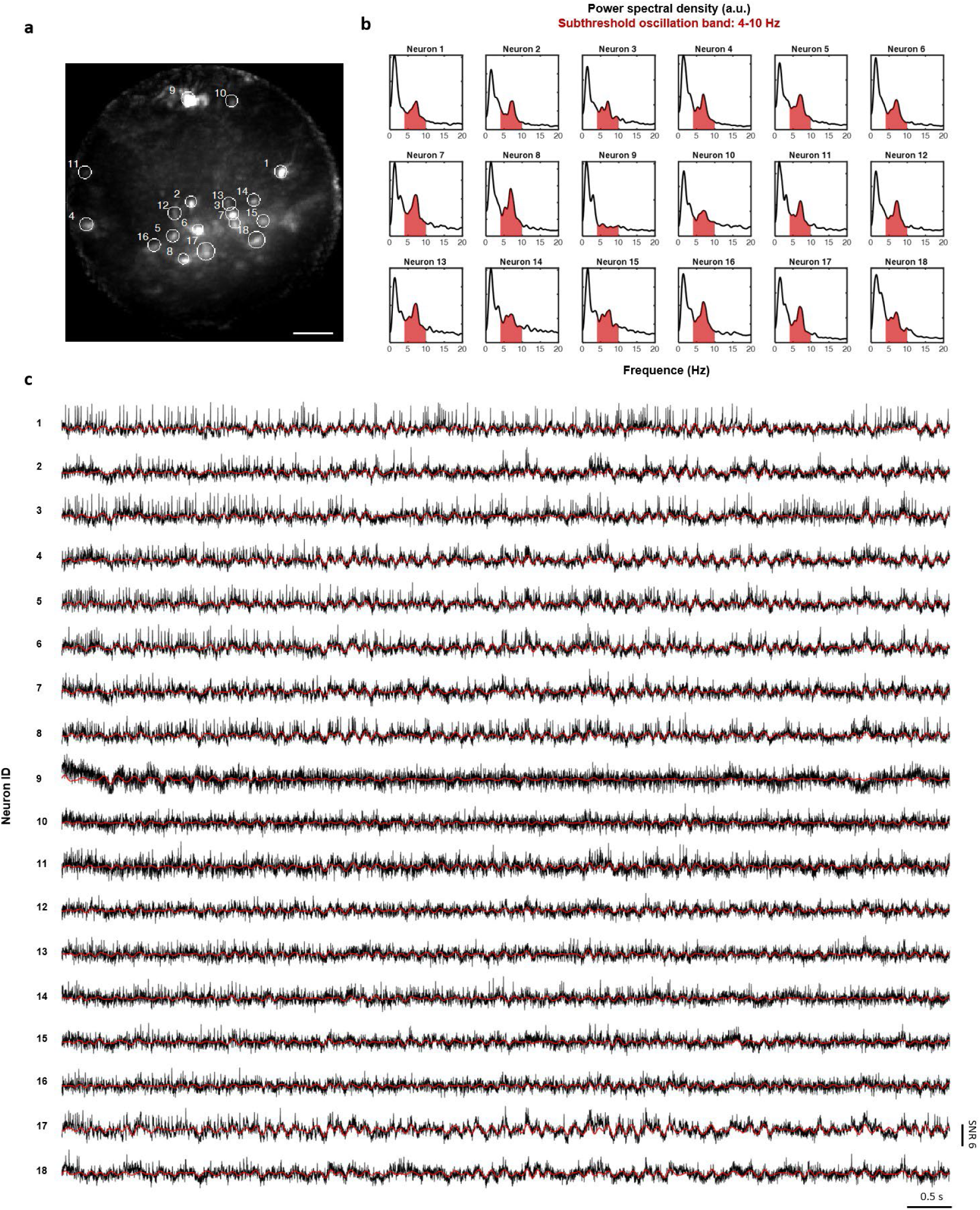
Observation of subthreshold oscillations in hippocampus interneurons. **a.** Example MIP of identical dataset used in **Fig. 4**, with neurons of interest labeled with number. Scale bar, 100 μm. **b.** The power spectral densities of 18 neurons traces throughout the entire recording (around three minutes). Red regions denote the frequency band known for theta oscillations. **c.** Raw traces of ten seconds with red lines plot the band-pass filtered signal in 4-10 Hz.

## References

1. Meng, G. et al. Ultrafast two-photon fluorescence imaging of cerebral blood circulation in the mouse brain in vivo. Proceedings of the National Academy of Sciences 119, e2117346119 (2022).

2. Abdelfattah, A. S. et al. Bright and photostable chemigenetic indicators for extended in vivo voltage imaging. Science 365, 699–704 (2019).

3. Abdelfattah, A. S. et al. Sensitivity optimization of a rhodopsin-based fluorescent voltage indicator. Neuron 111, 1547–1563.e9 (2023).

4. Hochbaum, D. R. et al. All-optical electrophysiology in mammalian neurons using engineered microbial rhodopsins. Nat Methods 11, 825–833 (2014).

5. Wu, J. et al. Kilohertz two-photon fluorescence microscopy imaging of neural activity in vivo. Nat Methods 17, 287–290 (2020).

6. Zhang, T. et al. Kilohertz two-photon brain imaging in awake mice. Nat Methods 16, 1119– 1122 (2019).

7. Wang, Z. et al. Imaging the voltage of neurons distributed across entire brains of larval zebrafish. 2023.12.15.571964 Preprint at 10.1101/2023.12.15.571964 (2023).

8. Weber, T. D., Moya, M. V., Kılıç, K., Mertz, J. & Economo, M. N. High-speed multiplane confocal microscopy for voltage imaging in densely labeled neuronal populations. Nat Neurosci 26, 1642–1650 (2023).

9. Sacconi, L. et al. KHz-rate volumetric voltage imaging of the whole Zebrafish heart. Biophysical Reports 2, 100046 (2022).

10. Voleti, V. et al. Real-time volumetric microscopy of in vivo dynamics and large-scale samples with SCAPE 2.0. Nat Methods 16, 1054–1062 (2019).

11. Pavani, S. R. P. et al. Three-dimensional, single-molecule fluorescence imaging beyond the diffraction limit by using a double-helix point spread function. Proceedings of the National Academy of Sciences 106, 2995–2999 (2009).

12. Llull, P. et al. Coded aperture compressive temporal imaging. Opt. Express, OE 21, 10526– 10545 (2013).

13. Wagadarikar, A. A., Pitsianis, N. P., Sun, X. & Brady, D. J. Video rate spectral imaging using a coded aperture snapshot spectral imager. Opt. Express, OE 17, 6368–6388 (2009).

14. Cui, Q., Park, J., Ma, Y. & Gao, L. Snapshot hyperspectral light field tomography. Optica, OPTICA 8, 1552–1558 (2021).

15. Prevedel, R. et al. Simultaneous whole-animal 3D imaging of neuronal activity using light-field microscopy. Nat Methods 11, 727–730 (2014).

16. Cong, L. et al. Rapid whole brain imaging of neural activity in freely behaving larval zebrafish (Danio rerio). eLife 6, e28158 (2017).

17. Skocek, O. et al. High-speed volumetric imaging of neuronal activity in freely moving rodents. Nat Methods 15, 429–432 (2018).

18. Zhang, Z. et al. Imaging volumetric dynamics at high speed in mouse and zebrafish brain with confocal light field microscopy. Nat Biotechnol 39, 74–83 (2021).

19. Wagner, N. et al. Instantaneous isotropic volumetric imaging of fast biological processes. Nat Methods 16, 497–500 (2019).

20. Wang, Z. et al. Real-time volumetric reconstruction of biological dynamics with light-field microscopy and deep learning. Nat Methods 18, 551–556 (2021).

21. Lu, Z. et al. Virtual-scanning light-field microscopy for robust snapshot high-resolution volumetric imaging. Nat Methods 20, 735–746 (2023).

22. Tian, T., Yuan, Y., Mitra, S., Gyongy, I. & Nolan, M. F. Single Photon Kilohertz Frame Rate Imaging of Neural Activity. Advanced Science 9, 2203018 (2022).

23. Guo, R., et al. EventLFM: Event Camera integrated Fourier Light Field Microscopy for Ultrafast 3D imaging. Preprint at 10.48550/arXiv.2310.00730 (2023).

24. Ashok, A. & Neifeld, M. A. Compressive light field imaging. in Three-Dimensional Imaging, Visualization, and Display 2010 and Display Technologies and Applications for Defense, Security, and Avionics IV vol. 7690 221–232 (SPIE, 2010).

25. Babacan, S. D. et al. Compressive Light Field Sensing. IEEE Transactions on Image Processing 21, 4746–4757 (2012).

26. Marwah, K., Wetzstein, G., Bando, Y. & Raskar, R. Compressive light field photography using overcomplete dictionaries and optimized projections. ACM Trans. Graph. 32, 46:1–46:12 (2013).

27. Antipa, N., Necula, S., Ng, R. & Waller, L. Single-shot diffuser-encoded light field imaging. in 2016 IEEE International Conference on Computational Photography (ICCP) 1–11 (2016). doi:10.1109/ICCPHOT.2016.7492880.

28. Liu, F. L., Kuo, G., Antipa, N., Yanny, K. & Waller, L. Fourier DiffuserScope: single-shot 3D Fourier light field microscopy with a diffuser. Opt. Express, OE 28, 28969–28986 (2020).

29. Yanny, K. et al. Miniscope3D: optimized single-shot miniature 3D fluorescence microscopy. Light Sci Appl 9, 171 (2020).

30. Antipa, N. et al. DiffuserCam: lensless single-exposure 3D imaging. Optica*, Vol.* 5*, Issue* *1**, pp.* 1-9 (2018) doi:10.1364/OPTICA.5.000001.

31. Feng, X., Ma, Y. & Gao, L. Compact light field photography towards versatile three-dimensional vision. Nat Commun 13, 3333 (2022).

32. Feng, X. & Gao, L. Ultrafast light field tomography for snapshot transient and non-line-of-sight imaging. Nat Commun 12, 2179 (2021).

33. Wang, Z., Hsiai, T. K. & Gao, L. Augmented light field tomography through parallel spectral encoding. Optica, OPTICA 10, 62–65 (2023).

34. Mandracchia, B. et al. High-speed optical imaging with sCMOS pixel reassignment. Nat Commun 15, 4598 (2024).

35. Guo, C. et al. Fourier light-field microscopy. Opt. Express, OE 27, 25573–25594 (2019).

36. Llavador, A., Sola-Pikabea, J., Saavedra, G., Javidi, B. & Martínez-Corral, M. Resolution improvements in integral microscopy with Fourier plane recording. Opt. Express, OE 24, 20792–20798 (2016).

37. Scrofani, G. et al. FIMic: design for ultimate 3D-integral microscopy of in-vivo biological samples. Biomed. Opt. Express, BOE 9, 335–346 (2018).

38. Levoy, M., Ng, R., Adams, A., Footer, M. & Horowitz, M. Light field microscopy. ACM Trans. Graph. 25, 924–934 (2006).

39. Broxton, M. et al. Wave optics theory and 3-D deconvolution for the light field microscope. Opt. Express, OE 21, 25418–25439 (2013).

40. Wang, Z. et al. A hybrid of light-field and light-sheet imaging to study myocardial function and intracardiac blood flow during zebrafish development. PLOS Computational Biology 17, e1009175 (2021).

41. Roustaei, M. et al. Computational simulations of the 4D micro-circulatory network in zebrafish tail amputation and regeneration. Journal of The Royal Society Interface 19, 20210898 (2022).

42. Zhou, Y., Zickus, V., Zammit, P., Taylor, J. M. & Harvey, A. R. High-speed extended-volume blood flow measurement using engineered point-spread function. Biomed. Opt. Express, BOE 9, 6444–6454 (2018).

43. Tomina, Y. & Wagenaar, D. A. A double-sided microscope to realize whole-ganglion imaging of membrane potential in the medicinal leech. eLife 6, e29839 (2017).

44. Briggman, K. L. & Kristan, W. B. Imaging Dedicated and Multifunctional Neural Circuits Generating Distinct Behaviors. J. Neurosci. 26, 10925–10933 (2006).

45. Briggman, K. L., Abarbanel, H. D. I. & Kristan, W. B. Optical Imaging of Neuronal Populations During Decision-Making. Science 307, 896–901 (2005).

46. Kannan, M. et al. Dual-polarity voltage imaging of the concurrent dynamics of multiple neuron types. Science 378, eabm8797 (2022).

47. Buzsáki, G. Theta Oscillations in the Hippocampus. Neuron 33, 325–340 (2002).

48. Gu, Z., et al. Hippocampal Interneuronal α7 nAChRs Modulate Theta Oscillations in Freely Moving Mice. Cell Reports 31, (2020).

49. Taxidis, J., Madruga, B., Melin, M. D., Lin, M. Z. & Golshani, P. Voltage imaging reveals that hippocampal interneurons tune memory-encoding pyramidal sequences. 2023.04.25.538286 Preprint at 10.1101/2023.04.25.538286 (2023).

50. Varga, C., Golshani, P. & Soltesz, I. Frequency-invariant temporal ordering of interneuronal discharges during hippocampal oscillations in awake mice. Proceedings of the National Academy of Sciences 109, E2726–E2734 (2012).

51. Yoon, Y.-G. et al. Sparse decomposition light-field microscopy for high speed imaging of neuronal activity. Optica, OPTICA 7, 1457–1468 (2020).

52. Vedula, V. et al. A method to quantify mechanobiologic forces during zebrafish cardiac development using 4-D light sheet imaging and computational modeling. PLOS Computational Biology 13, e1005828 (2017).

53. Wagner, N. et al. Deep learning-enhanced light-field imaging with continuous validation. Nat Methods 18, 557–563 (2021).

54. Zhang, Y. et al. DiLFM: an artifact-suppressed and noise-robust light-field microscopy through dictionary learning. Light Sci Appl 10, 152 (2021).

55. Pégard, N. C. et al. Compressive light-field microscopy for 3D neural activity recording. Optica, OPTICA 3, 517–524 (2016).

56. HiCAM: High-Speed, High-Sensitivity Imaging Camera - Lambert. lambertinstruments.com https://lambertinstruments.com/products/hicam (2020).

57. Lin, X., Wu, J., Zheng, G. & Dai, Q. Camera array based light field microscopy. Biomed. Opt. Express, BOE 6, 3179–3189 (2015).

58. Yi, C. et al. Video-rate 3D imaging of living cells using Fourier view-channel-depth light field microscopy. Commun Biol 6, 1–8 (2023).

59. Zhang, Y. et al. Computational optical sectioning with an incoherent multiscale scattering model for light-field microscopy. Nat Commun 12, 6391 (2021).

60. Zhang, Y. et al. Multi-focus light-field microscopy for high-speed large-volume imaging. PhotoniX 3, 30 (2022).

61. Zhu, L. et al. Sustained 3D isotropic imaging of subcellular dynamics using adaptive VCD light-field microscopy 2.0. 2023.03.15.532876 Preprint at 10.1101/2023.03.15.532876 (2023).

62. Biggs, D. S. C. & Andrews, M. Acceleration of iterative image restoration algorithms. Appl. Opt., AO 36, 1766–1775 (1997).

63. Ershov, D. et al. TrackMate 7: integrating state-of-the-art segmentation algorithms into tracking pipelines. Nat Methods 19, 829–832 (2022).

64. Tinevez, J.-Y. et al. TrackMate: An open and extensible platform for single-particle tracking. Methods 115, 80–90 (2017).

65. Tomina, Y. & Wagenaar, D. Dual-sided Voltage-sensitive Dye Imaging of Leech Ganglia. BIO-PROTOCOL 8, (2018).

66. Miller, E. W. et al. Optically monitoring voltage in neurons by photo-induced electron transfer through molecular wires. Proceedings of the National Academy of Sciences 109, 2114–2119 (2012).

67. Wetzel, A. W. et al. Registering large volume serial-section electron microscopy image sets for neural circuit reconstruction using FFT signal whitening. in 2016 IEEE Applied Imagery Pattern Recognition Workshop (AIPR) 1–10 (2016). doi:10.1109/AIPR.2016.8010595.

68. Thomson, D. J. Spectrum estimation and harmonic analysis. Proceedings of the IEEE 70, 1055–1096 (1982).

69. J, T. D. Jackknifed error estimates for spectra, coherences, and transfer functions. Advances in Spectrum Analysis and Arra Processing*. Vol.* 1 58–113 (1991).

70. Nichol, H., Amilhon, B., Manseau, F., Badrinarayanan, S. & Williams, S. Electrophysiological and Morphological Characterization of Chrna2 Cells in the Subiculum and CA1 of the Hippocampus: An Optogenetic Investigation. Front. Cell. Neurosci. 12, (2018).

71. Tarantino, N. et al. TNF and IL-1 exhibit distinct ubiquitin requirements for inducing NEMO– IKK supramolecular structures. Journal of Cell Biology 204, 231–245 (2014).

