## Supplementary Information for "Kilohertz volumetric imaging of in-vivo dynamics using squeezed light field microscopy"

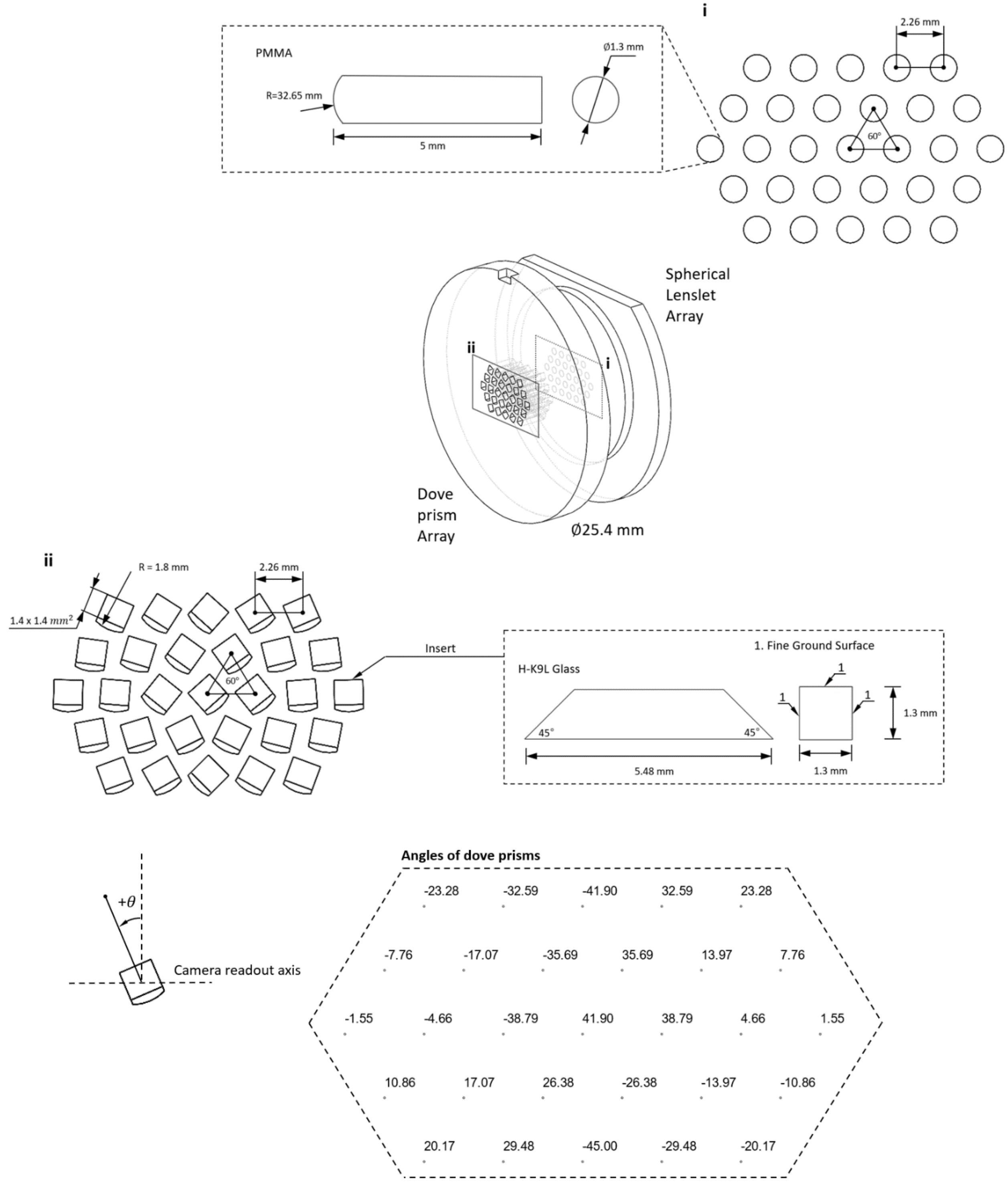


**Supplementary Figure 1. Design of lenslet and dove prism array.** (**i**) The lenslet array is in-house fabricated on a PMMA substrate. (**ii**) The holder of dove prims, the dove prism, and the angle arrangement of dove prisms.

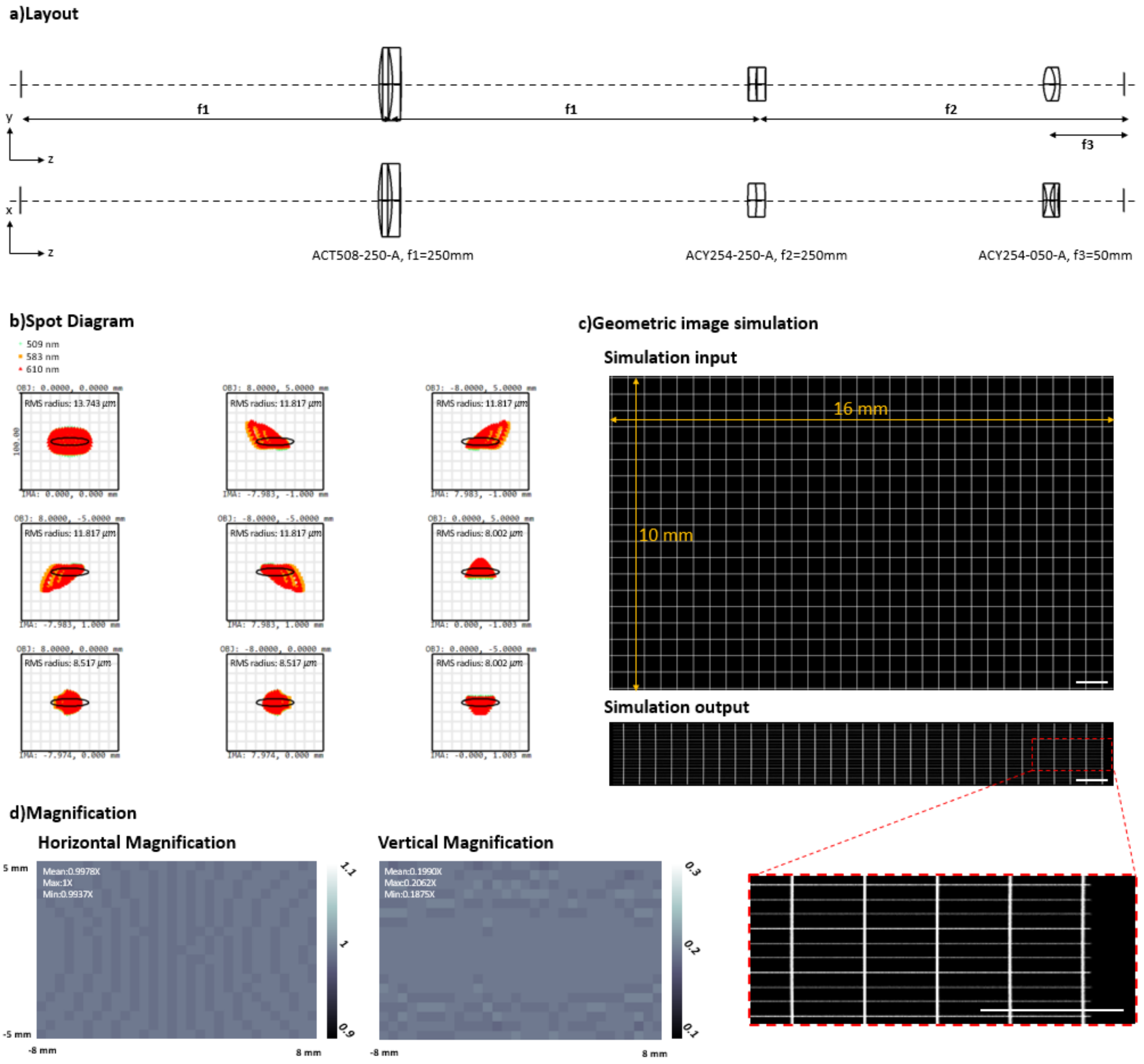


**Supplementary Figure 2. Design of anamorphic relay lens. a**. Layout of the relay system, consisting of one spherical achromatic doublet and two orthogonally placed cylindrical achromat doublets, where off-the-shelf components (Thorlabs) were used. **b**. Spot diagram shows aberration (within two times the pixel size, 6.5 $\mu m$) across the field of view of the lenslet array image (16 mm × 10 mm). **c**. OpticStudio Zemax simulations on an image of a grid. The red dotted box shows the zoom-in picture. Scale bar, 1 mm. **d**. Horizontal and vertical magnification measured from the grid simulation. The results show a uniform and constant scaling factor across the field of view, which ensures correct image transformation for all sub-aperture images.

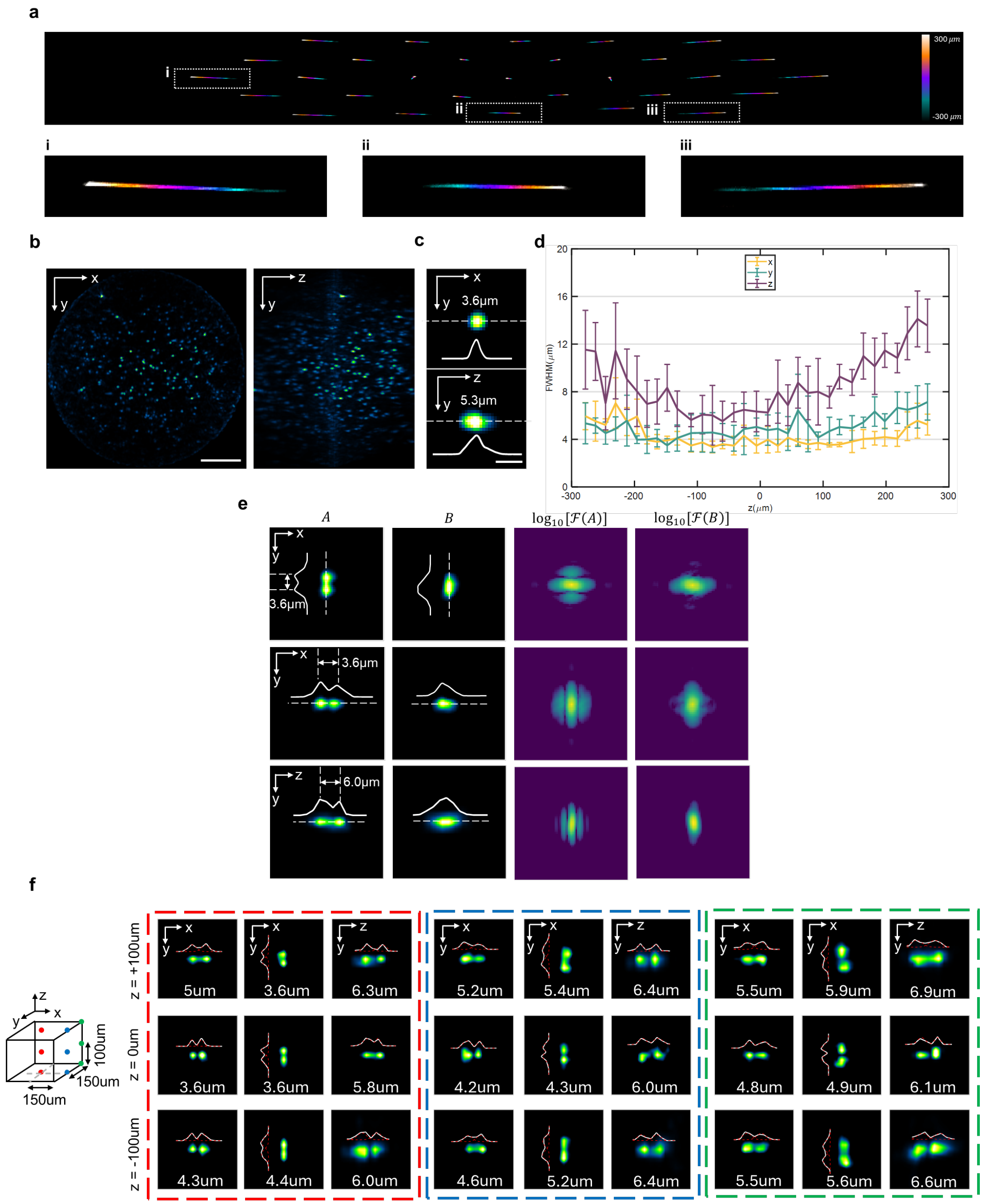


**Supplementary Figure 3. Characterization of lateral and axial resolutions of SLIM. a**. x-y MIP of the SLIM PSFs with depth color-coded. **b**. fluorescent beads. Scale bar 100 µm. **c**. Individual reconstructed bead in (a) in lateral(x-y) and axial(y-z) dimensions and the corresponding profiles along the dashed lines. Scale bar, 5 µm. **d**. Average axial (z) and lateral (x,y) FWHM of the beads across the volumes reconstructed by SLIM. Center lines represent means and error bars denote standard deviations. **e**. Analysis of the reconstructed images of two virtually separated beads obtained by SLIM with cross-section profiles along the dashed lines. We imaged the same 1µm bead at two positions, adjusting the interval gradually using a piezo translation stage. By combining the images obtained from these two positions, we created two virtually separated beads with an arbitrary distance. The first and second columns (A and B) are the reconstructed images of resolved beads (A) and unresolved beads (B). The third and fourth columns are the Fourier analysis of the first and second columns using function: f(x)=log(|F(x)|), where F(x) represents the Fourier transform. **f**. Two virtually separated beads resolution measurements at several different places within the volume. At each lateral location (red, blue, green box), the experiments are repeated at different axial locations (-100, 0, 100 µm). The beads are displaced horizontally, vertically and axially.

**a**. Selective volume side-illumination setup

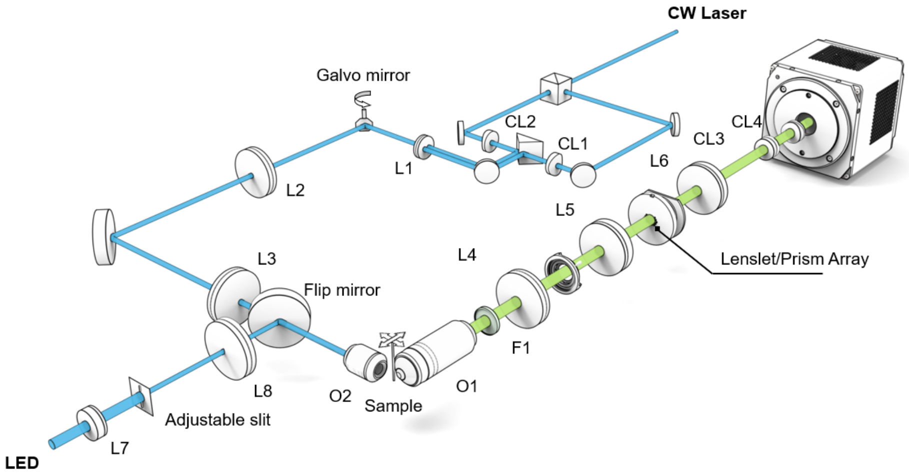


**b**. Widefield epi-illumination setup

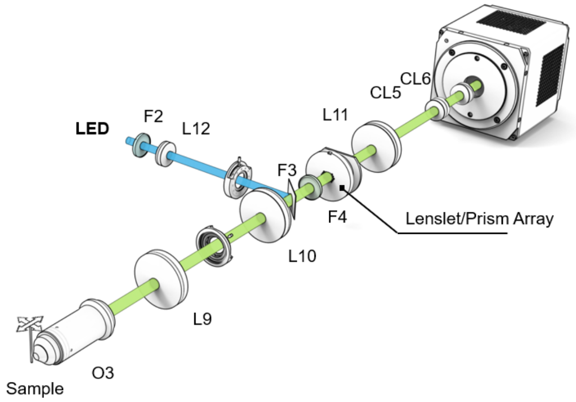


**Supplementary Figure 4. Schematics of the SLIM with different illumination systems.** L1-L12, lens; CL1-CL6, cylindrical lens. O1-O3, objectives. F1-F4, filters. See **Supplementary Table 3** for details.

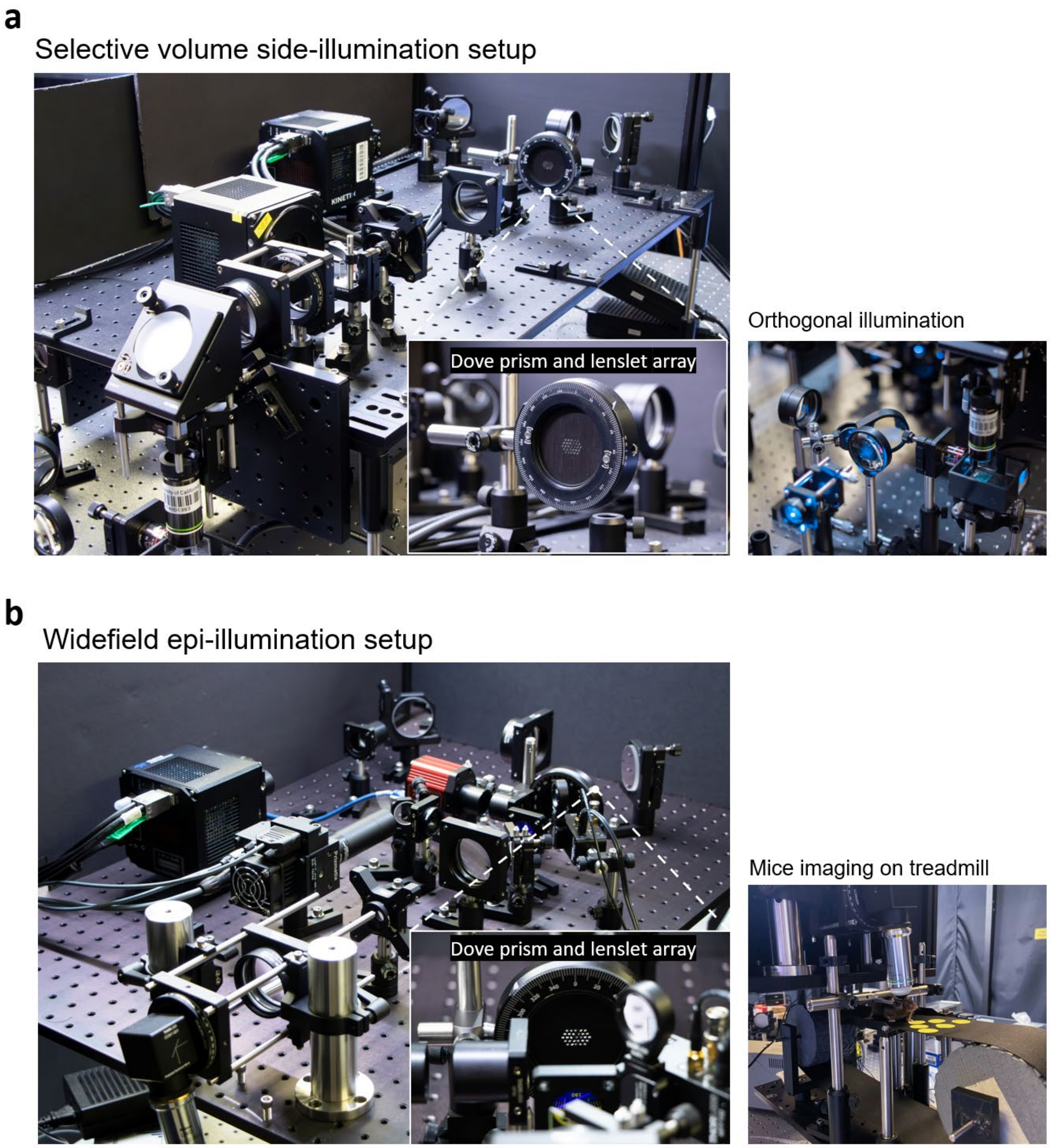


**Supplementary Figure 5. Photographs of the setups.**

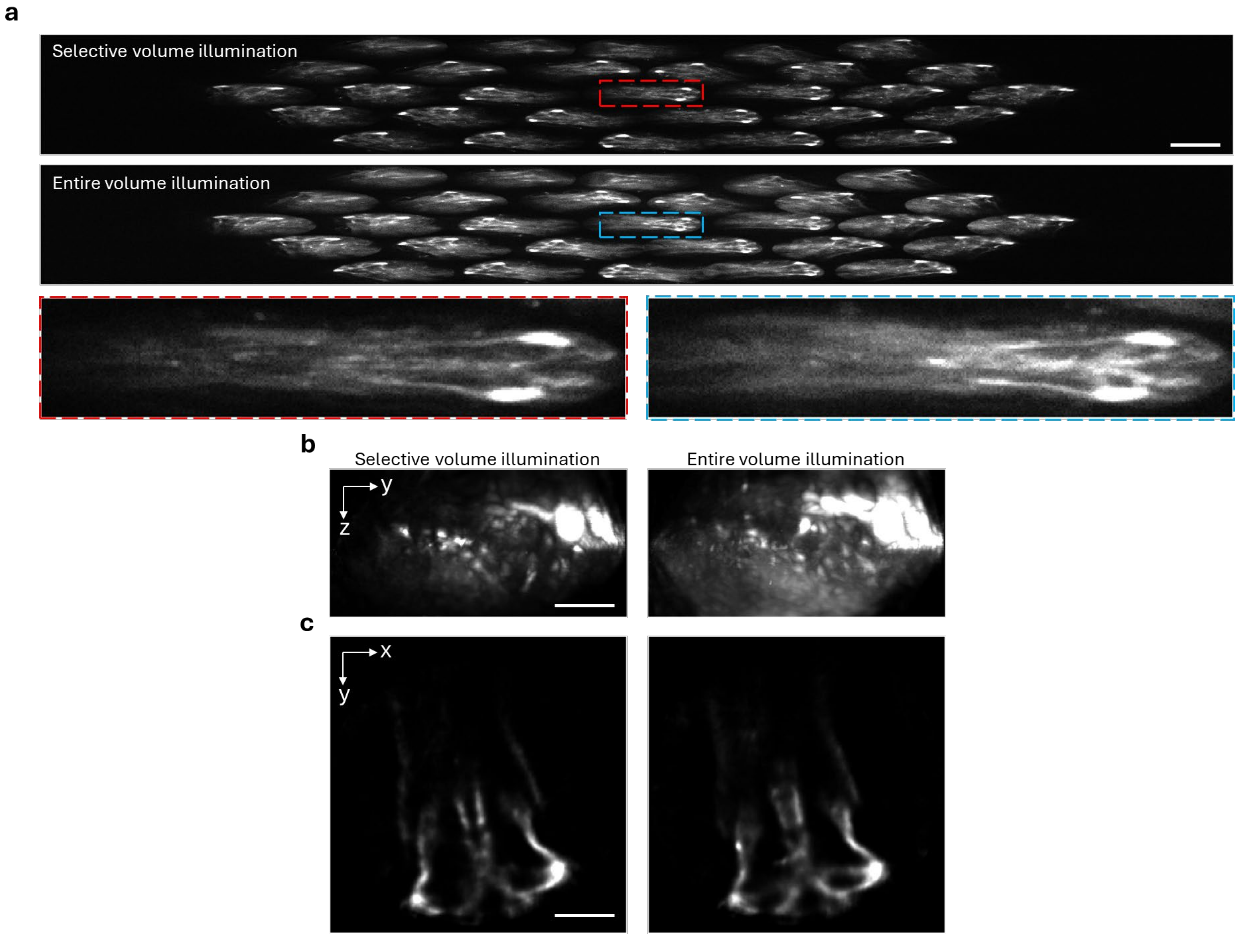


**Supplementary Figure 6. Comparisons of image quality of SLIM under different illumination configuration in zebrafish brain. a.** Raw measurement of embryonic zebrafish brain under selective volume illumination. Selective volume illumination (1$^{st}$ row) was applied to suppress fluorescence outside the imaging volume. The entire volume is illuminated (2$^{nd}$ row). The red and blue dashed box indicate the zoom in view of center perspective. Scale bar, 250 µm. **b.** MIP of SLIM reconstruction for zebrafish brain. Scale bar, 100 µm. **c.** The representative slice of SLIM reconstruction for zebrafish brain. Scale bar, 100 µm. Left: selective volume illumination. Right: entire volume illumination.

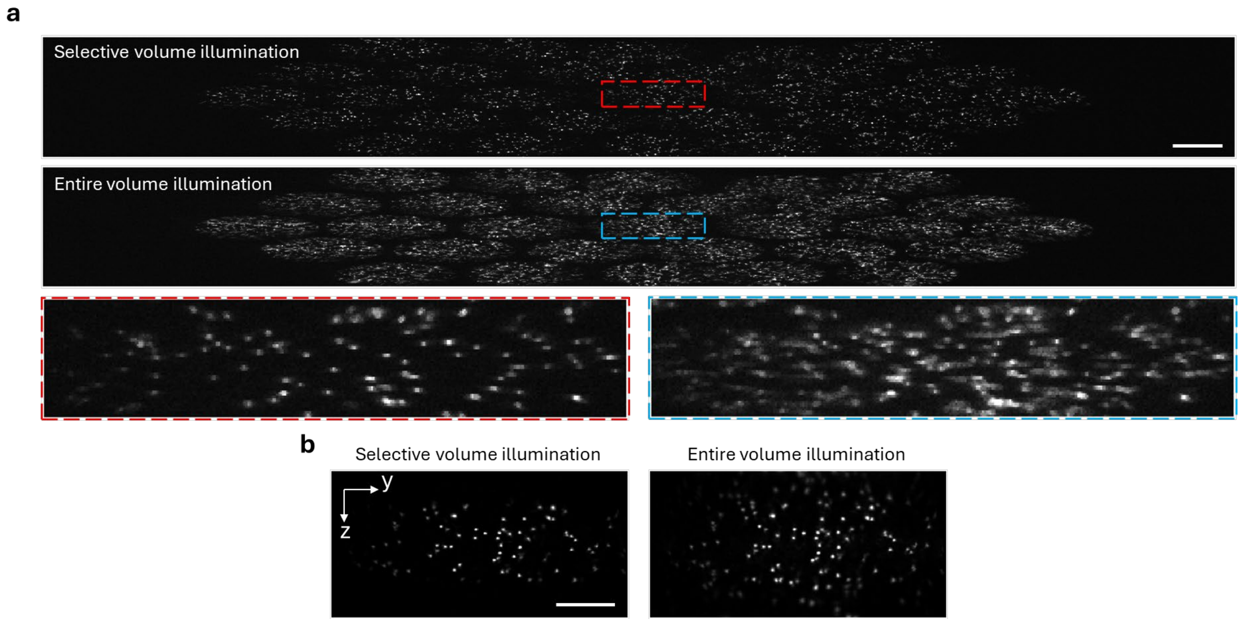


**Supplementary Figure 7. Comparisons of image quality of SLIM under different illumination configuration in fluorescent beads. a.** Raw measurement of fluorescent beads under selective volume illumination. Selective volume illumination (1$^{st}$ row) was applied to suppress fluorescence outside the imaging volume. The entire volume is illuminated (2$^{nd}$ row). The red and blue dashed box indicate the zoom in view of center perspective. Scale bar, 250 µm. **b.** MIP of SLIM reconstruction of fluorescent beads. Scale bar, 100 µm.

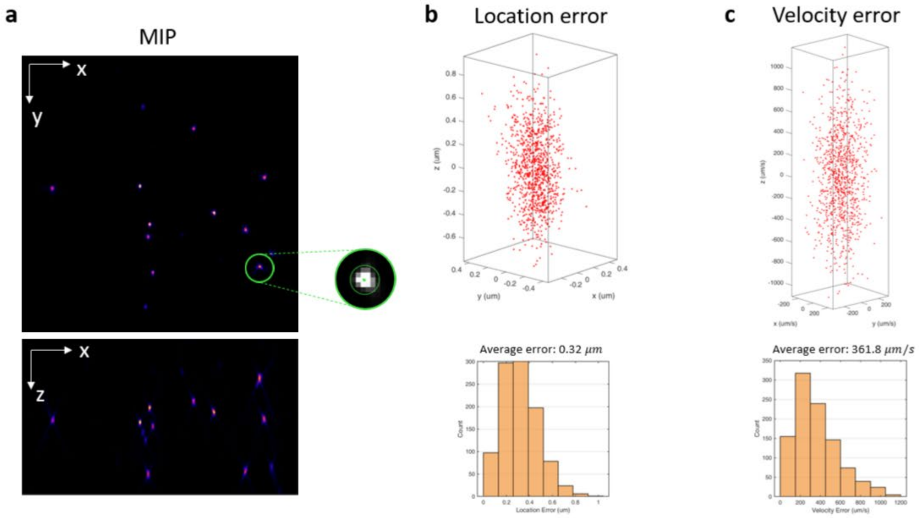


**Supplementary Figure 8. Tracking errors quantified by imaging static fluorescent beads at 1,000 vps. a.** x-y and x-z MIPs of the reconstruction of the example frame. Beads are assumed to be stationary, and the tracking error is indicated by the tracking difference among different frames. These sub-pixel errors potentially come from the reconstruction and tracking algorithms and environment vibration. **b.** Location variation. Each red dot plots the relative position of the tracking position in one frame to the average position across all frames (n=1000). The histogram plots the length of the location error vector. **c.** Velocity error caused by the location variation in adjacent frames. The histogram plots the absolute value of the velocity vector.

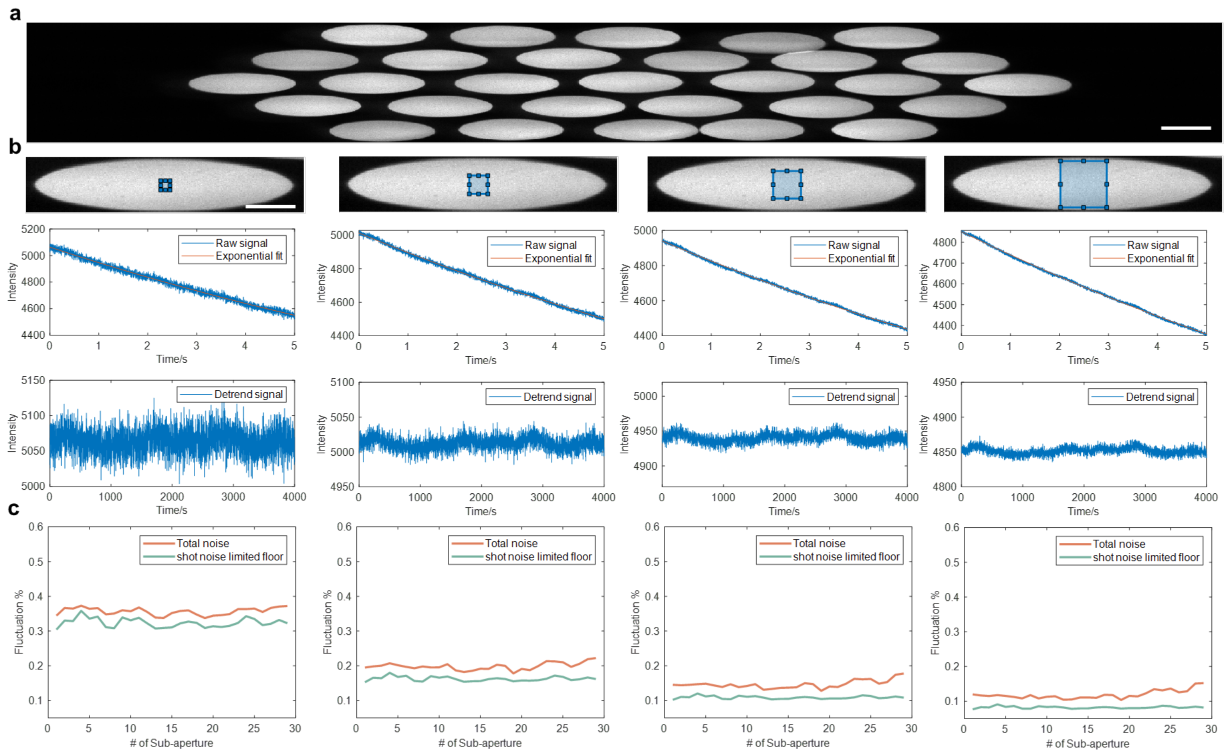


**Supplementary Figure 9. Noise characterization in ultra-low-noise LED illumination a.** Raw image of reference fluorescence dye. Scale bar 200 µm. **b.** Central sub-aperture images with various region of interest (10*10, 20*20, 30*30, 50*50), and the corresponding averaged time series signal. Scale bar, 100 µm. We assume that the signal of fluorescence dye is uniform across the entire field of view. Photobleaching is observed and detrended by exponential fitting. Both total signal noise $Mean(signal)/std(signal)$ and shot noise $\sqrt{number\ of\ pixel}/\sqrt{Mean(signal)}$ is calculate for different size of ROI. (0.35%/0.31%, 0.19%/0.16%, 0.13%/0.10%, 0.09%/0.064%) **c.** All 29 sub-aperture images with various region of interest (10*10, 20*20, 30*30, 50*50), and the corresponding fluctuation percentage.

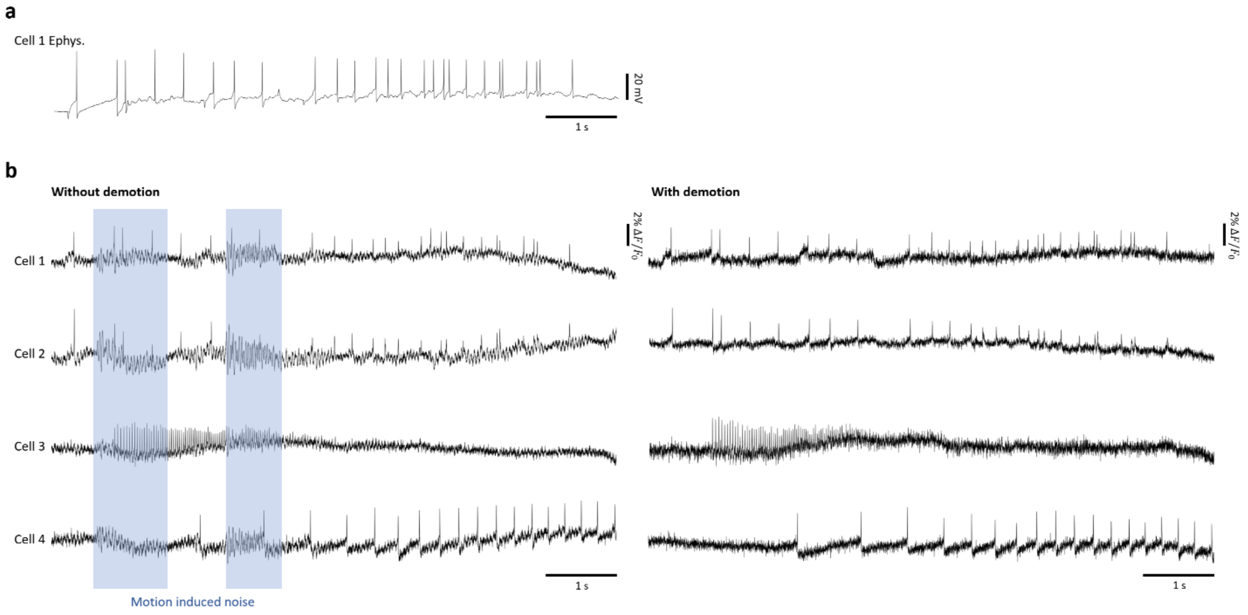


**Supplementary Figure 10. Image demotion for optical recording of membrane action potentials in leech. a.** The ground-truth electrophysiological signal provided by electrode on cell 1. **b**. By assuming the sample being static during recording, motion correction has been applied to the image sequence by image registration between adjacent frames. With motion correction, the noise induced by sample/environment vibration can be suppressed. The blue boxes label the example time window when such noise appears severe and affects the detection of voltage spikes.

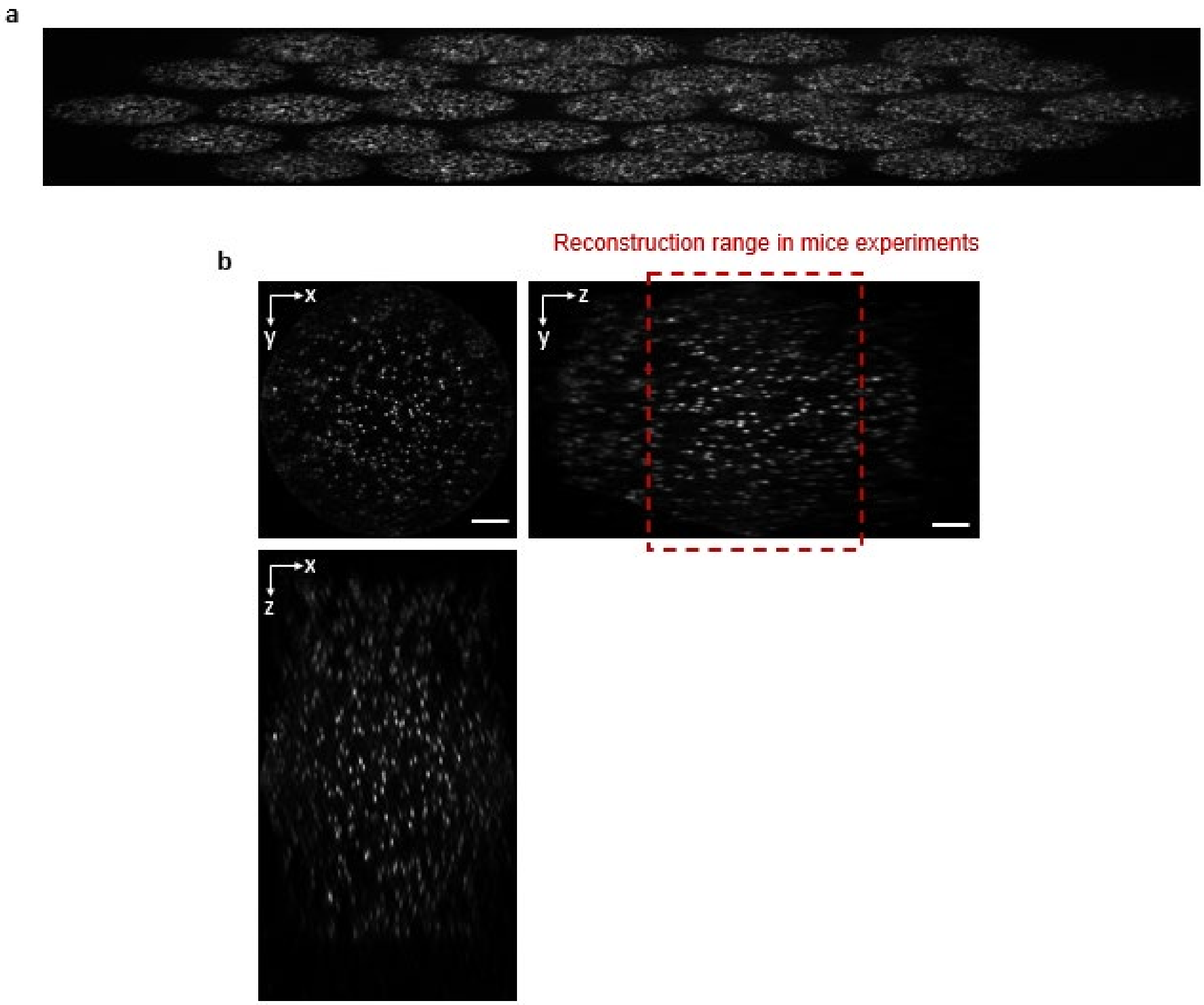


**Supplementary Figure 11. Fluorescent beads under mouse imaging setup. a.** Raw measurement. ROI is configured to be 330 × 2400 pixels to fit 29 sub-aperture images (each 61 × 305 pixels). **b**. The reconstruction result (305 × 305 × 151 pixels, FOV ∅ 688 × 1204 µm) of measurement in **a**. Inset, the average pixel intensity per depth. The red dotted box indicated the depth range (-296 ~ 296 µm) used in mouse imaging experiments. Scale bar, 100 µm.

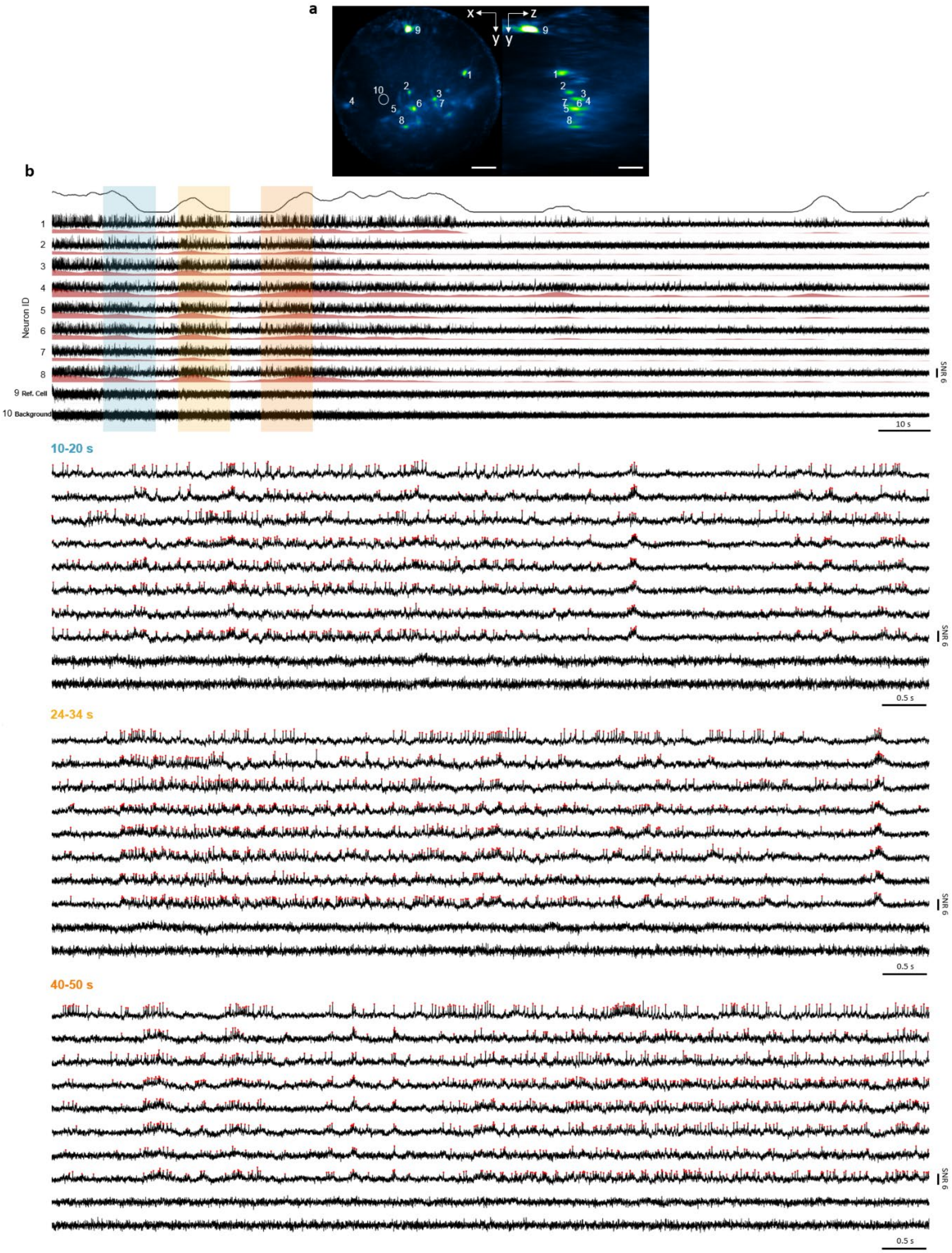


**Supplementary Figure 12. Zoom-in views of different time windows of data presented in Fig. 4. Traces 9 and 10 are an inactive neuron and a background measurement, respectively, to confirm the fidelity of other traces.**

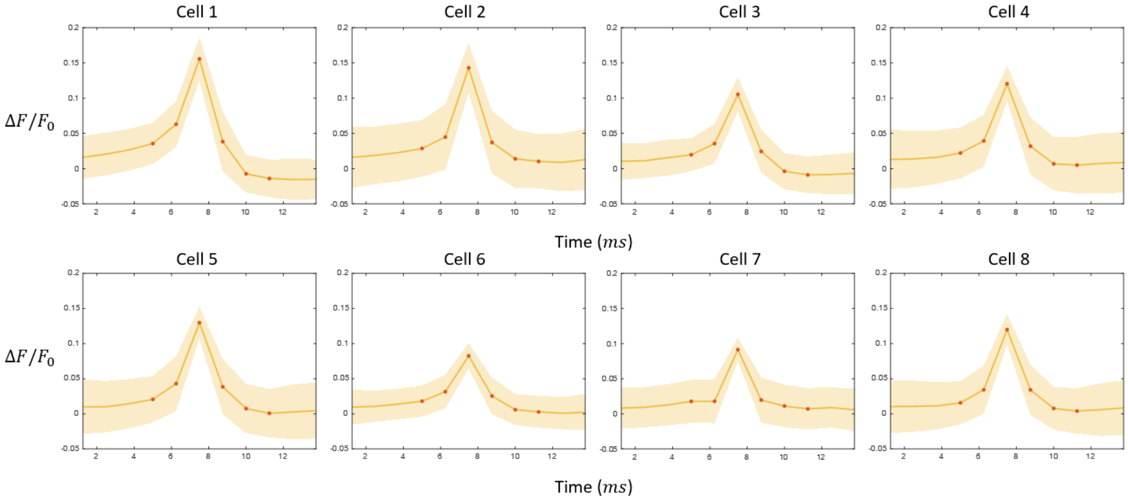


**Supplementary Figure 13. Average spike waveforms for neurons demonstrated in Fig. 4.** Interval between sampling points (orange) is 1.25 ms.

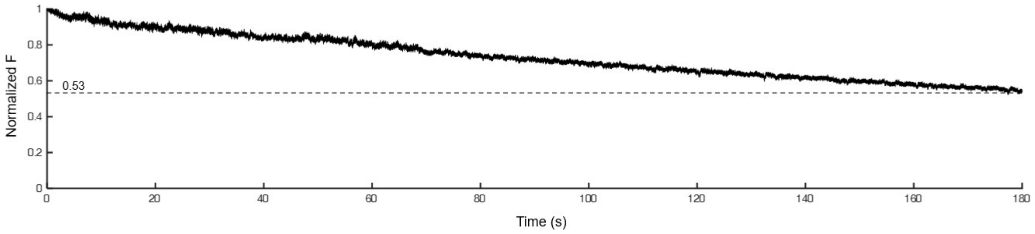


**Supplementary Figure 14. Representative pAce fluorescence photobleaching during 180s continuous SLIM imaging.** The mean pixel intensity for neurons (n=25) in the FOV in **Supplementary figure 12** The illumination power on sample is measured to be 36mW/mm$^2$.

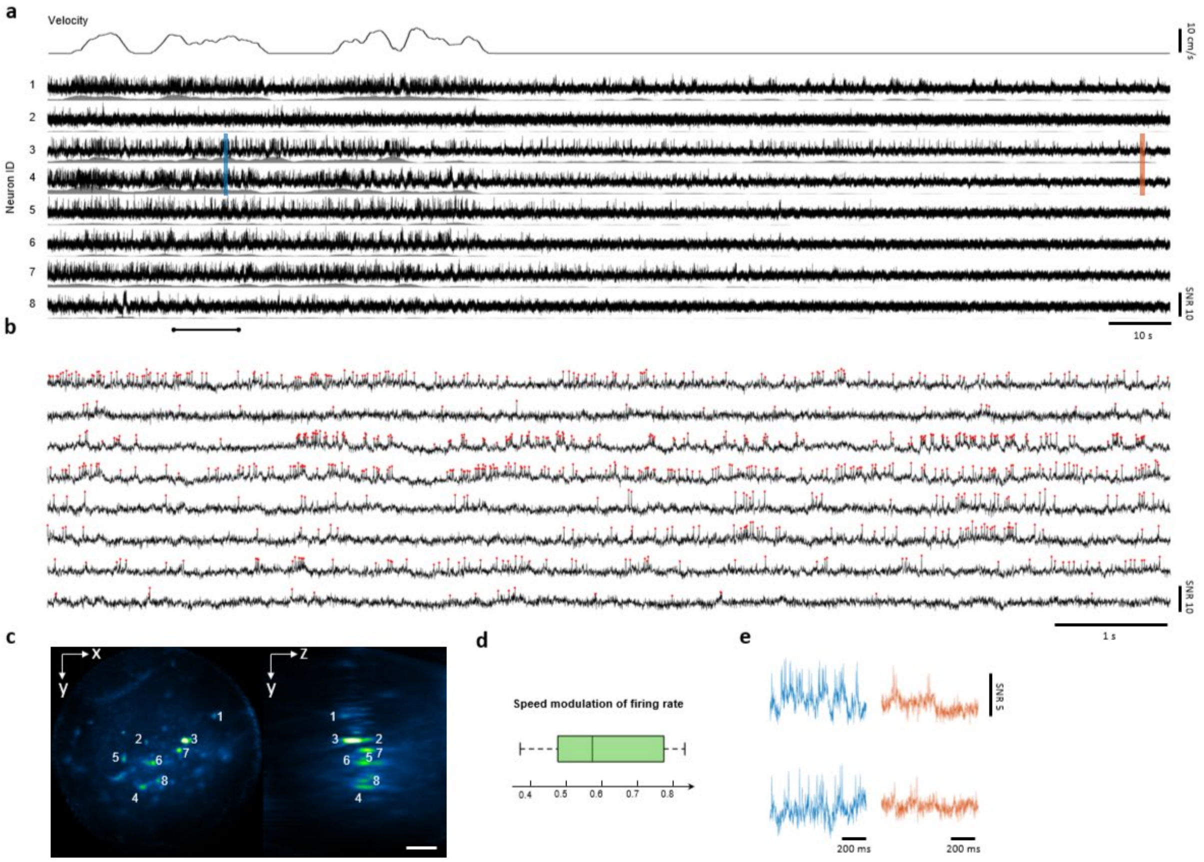


**Supplementary Figure 15. 3D voltage imaging of hippocampus in behaving mice at 800 vps.** Another example results of continuous imaging across 180s while animal runs on treadmill, verifying the findings on speed modulation of neuron firing rate in **Fig. 4**. **a.** Detrended fluorescent signal traces with firing rate. **b.** The zooms in on the window (20 – 30s) in **a**.The red dots denote the detected spikes. **c.** 3D MIP of SLIM reconstruction. Scale bar, 100 µm. **d.** Pearson correlation coefficients between firing rate and animal velocity. **e.** Representative signal traces in the blue and orange windows in **a.**

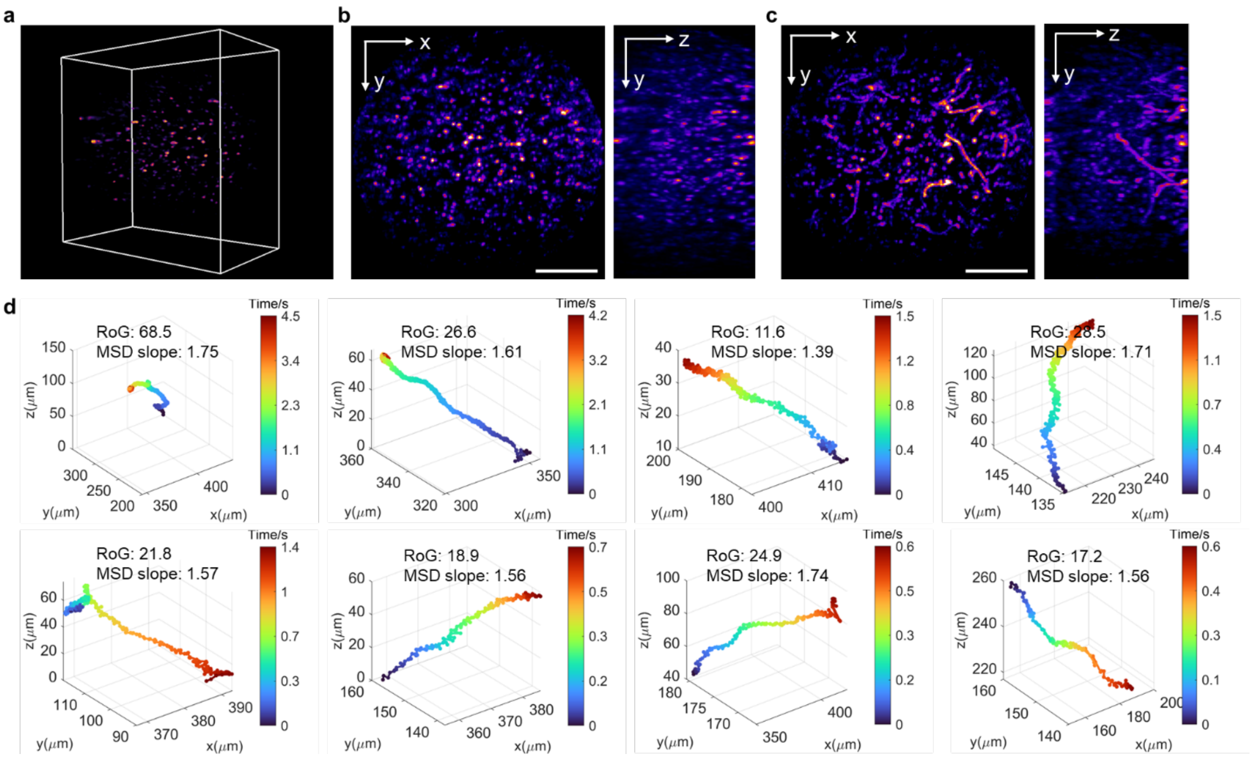


**Supplementary Figure 16. 3D imaging of free-swimming *Vibrio cholerae* bacteria at 200 vps. a.** 3D rendering volume of *Vibrio cholerae* bacteria. **b.** MIP from x-y and y-z slices of *Vibrio cholerae* bacteria. *Vibrio cholerae* are stained with an external membrane stain. The total recording time was 5s. **c.** MIPs of the swimming bacteria trajectory obtained by combining frames over time. **d.** Representative trajectories of swimming bacteria with their respective Radius of Gyration (RoG) and Mean Squared Displacement (MSD) slope measurements labeled. Scale bar, 120 µm.

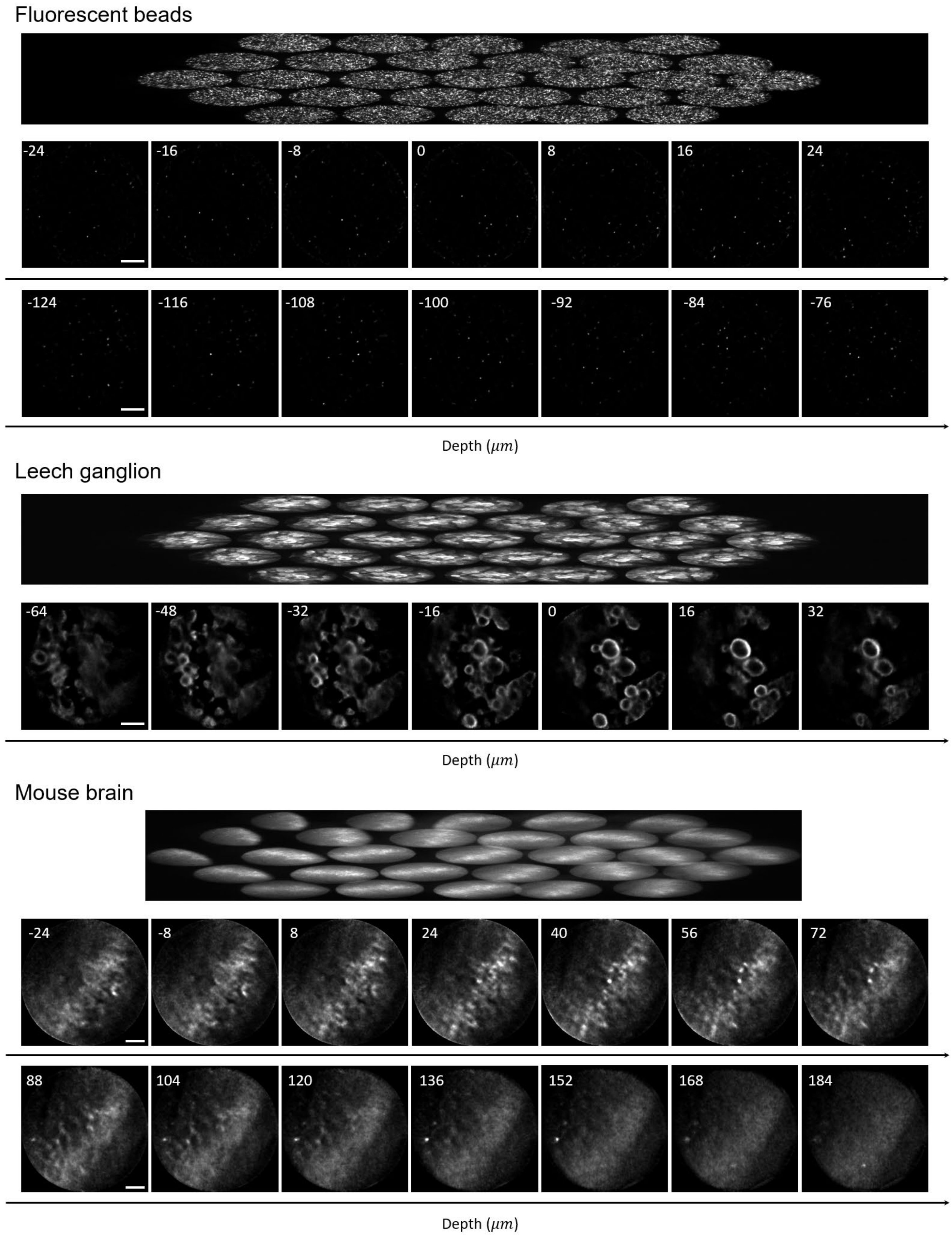


**Supplementary Figure 17. Raw and reconstruction slices of representative samples.**

**Supplementary Note 1**

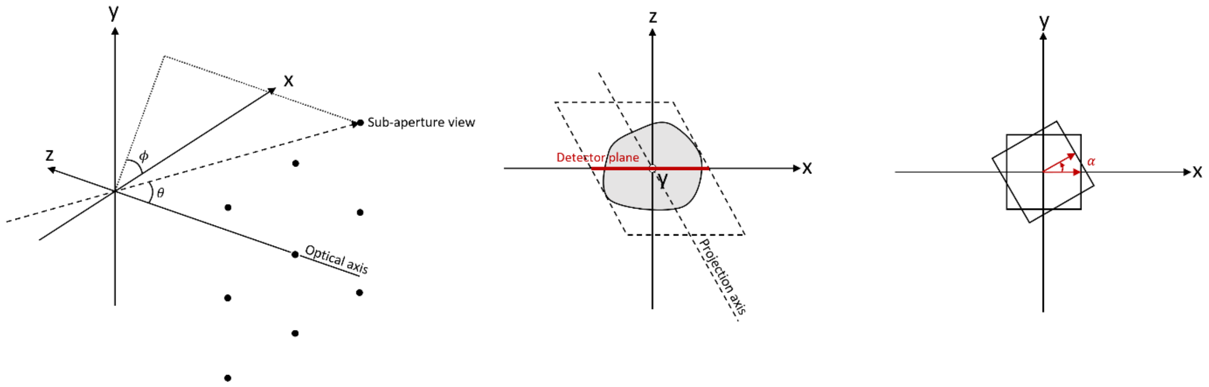


In this note, we model each sub-aperture image in SLIM/Fourier LFM as a parallel projection along the line of sight at the sub-aperture's view angle $\theta$ and $\phi$. However, in contrast to classical tomographic system where detector rotates with scanner, there's an oblique angle between projection axis and detector plane. Following Fourier slice theorem, we attempt to find the mapping between the spectrum of 2D SLIM image and the 3D spectrum of the object.

For a 3D distributed signal $f(x, y, z)$, the 2D sub-aperture view $I(x, y)$ is:

$$I(x, y) = \int_z f(x - z\ tan\theta\ cos\phi, y - z\ tan\theta\ sin\phi, z)\, dz$$

Meanwhile, the coordinate transformation introduced by the in-plane rotation is:

$$\begin{bmatrix} x' \\ y' \end{bmatrix} = \begin{bmatrix} cos\alpha & -sin\alpha \\ sin\alpha & cos\alpha \end{bmatrix} \begin{bmatrix} x \\ y \end{bmatrix}$$

$$f'(x', y', z') = f(x, y, z) = f(x'\ cos\alpha + y'\ sin\alpha, -x'\ sin\alpha + y'\ cos\alpha, z')$$

The 2D Fourier transformation of the in-plane rotated projection $I'(x', y')$ is:

$$\begin{aligned} \mathcal{F}\{I'(x', y')\} &= \iint_{x', y'} I'(x', y')\, e^{-i2\pi(u'x'+v'y')}\, dx'\, dy' \\ &= \iint_{x', y'} \int_{z'} f'(x' - z'\ tan\theta\ cos\phi, y' - z'\ tan\theta\ sin\phi, z')\, dz'\, e^{-i2\pi(u'x'+v'y')}\, dx'\, dy' \\ &= \iint_{x', y'} \int_{z} f(x'\ cos\alpha + y'\ sin\alpha - z'\ tan\theta\ cos\phi, -x'\ sin\alpha + y'\ cos\alpha - z'\ tan\theta\ sin\phi, z')\, dz' \\ &\qquad e^{-i2\pi(u'x'+v'y')}\, dx'\, dy' \end{aligned}$$

Let

$$
\begin{aligned}
x &= x' \, cos\alpha + y' \, sin\alpha - z' \, tan\theta \, cos\phi \\
y &= -x' \, sin\alpha + y' \, cos\alpha - z' \, tan\theta \, sin\phi \\
z &= z'
\end{aligned}
$$

Then

$$
\begin{aligned}
x' &= xcos\alpha - ysin\alpha + ztan\theta cos(\alpha + \phi) \\
y' &= xsin\alpha + ycos\alpha + ztan\theta sin(\alpha + \phi) \\
z' &= z
\end{aligned}
$$

The Jacobian of the $x, y$ coordinate transformation is:

$$
J(x, y) = \begin{vmatrix} \frac{\partial x'}{\partial x} & \frac{\partial x'}{\partial y} \\ \frac{\partial y'}{\partial x} & \frac{\partial y'}{\partial y} \end{vmatrix} = \begin{vmatrix} cos\alpha & -sin\alpha \\ sin\alpha & cos\alpha \end{vmatrix} = 1
$$

Therefore, we can rewrite the 2D Fourier transformation as:

$$
\begin{aligned}
\mathcal{F}\{I'(x', y')\} &= \iint_{x,y} \int_z f(x, y, z) \, dz \\
& e^{-i2\pi\{u'xcos\alpha - u'ysin\alpha + u'ztan\theta cos(\alpha+\phi) + v'xsin\alpha + v'ycos\alpha + v'ztan\theta sin(\alpha+\phi)\}} J(x, y) \, dx \, dy \\
&= \iiint_{x,y,z} f(x, y, z) e^{-i2\pi\{u'xcos\alpha - u'ysin\alpha + u'ztan\theta cos(\alpha+\phi) + v'xsin\alpha + v'ycos\alpha + v'ztan\theta sin(\alpha+\phi)\}} \, dx \, dy \, dz \\
&= \iiint_{x,y,z} f(x, y, z) e^{-i2\pi\{ux + vy + wz\}} \, dx \, dy \, dz
\end{aligned}
$$

where

$$
\begin{aligned}
u &= u'cos\alpha + v'sin\alpha \\
v &= -u'sin\alpha + v'cos\alpha \\
w &= u'tan\theta cos(\alpha + \phi) + v'tan\theta sin(\alpha + \phi)
\end{aligned}
$$

Note that it has the same form of 3D Fourier transformation of $f(x, y, z)$:

$$
\mathcal{F}\{f(x, y, z)\} = \iiint_{x,y,z} f(x, y, z) e^{-i2\pi\{ux + vy + wz\}} \, dx \, dy \, dz
$$

With the coordinate transformation between $u, v, w$ and $u', v', w'$ we can map the 2D spectrum of projection $I'(x', y')$ to the 3D Fourier space of $f(x, y, z)$ and reveal the 3D frequency components that are retrievable from the 2D measurement. We assume that the squeezed measurement undersamples the sub-aperture image in one dimension and thus delivers an elliptical transfer function (with long axis in $u'$ and short axis in $v'$ direction). According to the coordinate transformation, we would pursue $\alpha + \phi$ to be zero in order to maximize the axial cutoff frequency, which indicates that the in-plane rotation angle should be chosen in attempt to align the disparity shift horizontally ($x'$ axis, camera rows). For example, the following figure shows how the aforementioned coordinate transformation applies to a sub-aperture view in our design ($\theta = 24.35°, \phi = -60°, \alpha = 58.97°$). The 2D elliptical spectrum pursues the $w$ direction in the 3D Fourier space with its long axis.

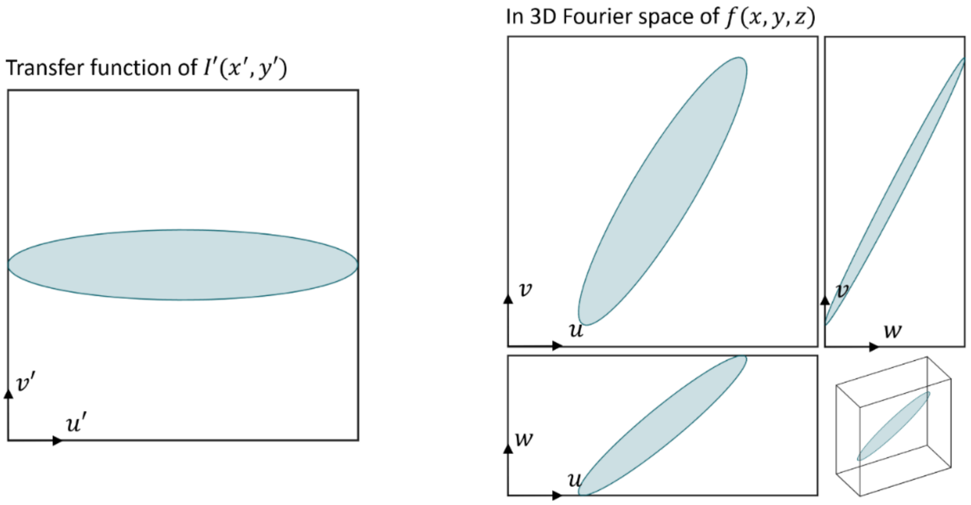


The effect of different strategies in angle selection can also be visualized in space domain. The figure below shows x-y MIP of simulated SLIM PSF. By examining the lateral shift induced by axial translation, we can infer the axial resolving capability. Strategy 1 allows the lateral shift to align with camera rows where we have full digital sampling power, in contrast to strategy 2 that does the opposite. We should note that these selection strategies cannot be exactly achieved since the entire angle set is uniformly distributed in 180 degree.

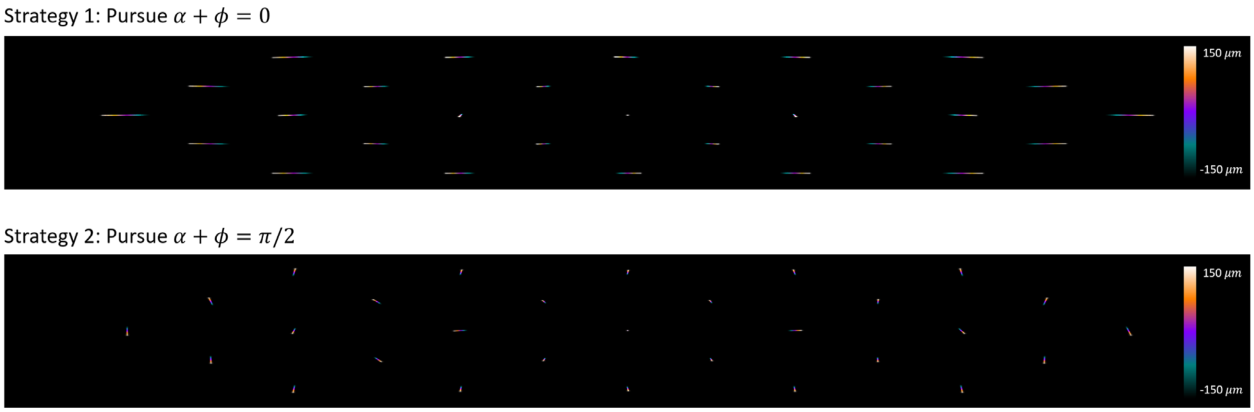

**Supplementary Note 2**

In this note, we present a series of simulations designed to evaluate the limitation and trade-off of our proposed SLIM system. It examines the illness of reconstruction and the assumption on sample signal sparsity. It also compares SLIMs against Fourier LFM (FLFM) and between various design parameters including number of sub-aperture images and vertical scaling ratios.

The 3D PSF of FLFM was first simulated following the method presented in Guo, C, et al.[1]. It was then cropped based on the lenslet position, with each patch being the 3D PSF of the sub-aperture. After calculating and assigning optimal angles (**Supplementary Note 1**), we applied in-plane rotation and then vertical scaling to obtain SLIM sub-aperture PSFs.

To synthesize corresponding SLIM measurement, we first collected ground truth (GT) 3D images. For synthetic zebrafish brain neurons, high-resolution image stack and neuron segmentation were acquired from Vladimirov N, et al. [2]. And following Cong, L, et al.[3], we randomly sampled a portion of neurons. From about 80,000 neurons we sampled 5%, 10%, 15% and 20%, which gave us n=3,585, 7,052, 10,565 and 21,358 number of neurons in our FOV. For real biological samples, we directly downloaded dataset for embryonic zebrafish vascular structure[4] and mouse brain neurons[5,6] and cropped/zero-padded it to fit our FOV. Next, for each SLIM sub-aperture PSF, the ground truth 3D image went through the corresponding rotation and vertical scaling. A depth-by-depth 2D convolution was performed between the transformed ground truth image and PSF. A depth-wise summation and adding Poisson noise gave the sub-aperture measurement. After repeating for every SLIM sub-aperture, the synthetic measurement was in form of a stack of sub-aperture images. We used Richardson-Lucy deconvolution for reconstruction.

**1.Sample sparsity and reconstruction illness**

SLIM is a compressive detection strategy for light field microscopy and its successful reconstruction relies on the sparseness of the sample, which we assume applies to a wide range of high-speed biological dynamics. As shown in **Supplementary Note 2 Fig. 1**, SLIM's performance can approximate FLFM when neurons are sparsely distributed (n=3,585, 7,052). However, it will degrade with higher density (e.g. n=21,358) and reconstructed neurons can no longer be clearly separated as density continues to increase. It's worth noting that SLIM holds up well in the sparseness settings used in FLFM literatures, where only 10% of neurons are assumed to be active at a given time[3,7].
A similar trend is expected when dealing with real sample datasets. From **Supplementary Note 2 Fig. 2**, SLIM shows comparable spatial resolution to FLFM on sparse zebrafish vasculature and mouse neurons. However, as shown in 5th row, when sample density and structural complexity increase significantly, SLIM's reconstruction quality begins to degrade. We note that SLIM is not intended for high-resolution structural imaging of highly complex samples, but these simulations underscore its robustness in scenarios where spatiotemporal sparsity is present.

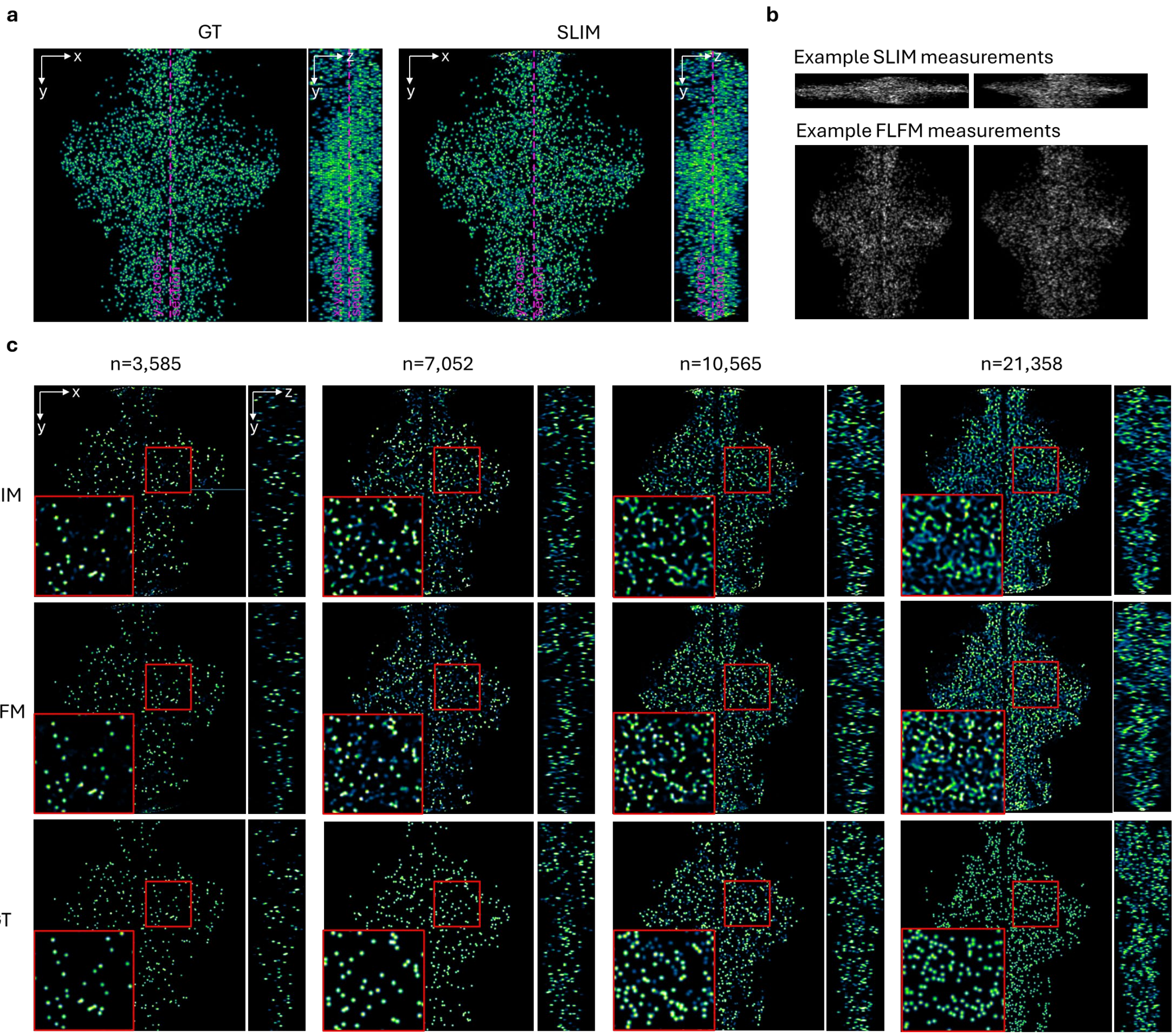


**Supplementary Note 2 Fig. 1. Simulation results of synthetic neurons in zebrafish with different density of neurons. a.** 3D MIP of synthetic neurons (GT) and corresponding SLIM reconstruction. **b.** Synthesized measurements of SLIM (top) and FLFM (bottom). Simulation adds Poison noise. **c.** Comparison between SLIM and FLFM reconstruction results with different density of neurons. Cross-sectioning slices are sampled at the position indicated by purple dotted line in **a**. From left to right, the ratio of randomly sampled neurons (83,890 in total) increases from 5%, 10%, 15% to 20%. The corresponding neuron number in our FOV is 3,585, 7,052, 10,565 and 21,358. The red box provides the zoom in view.

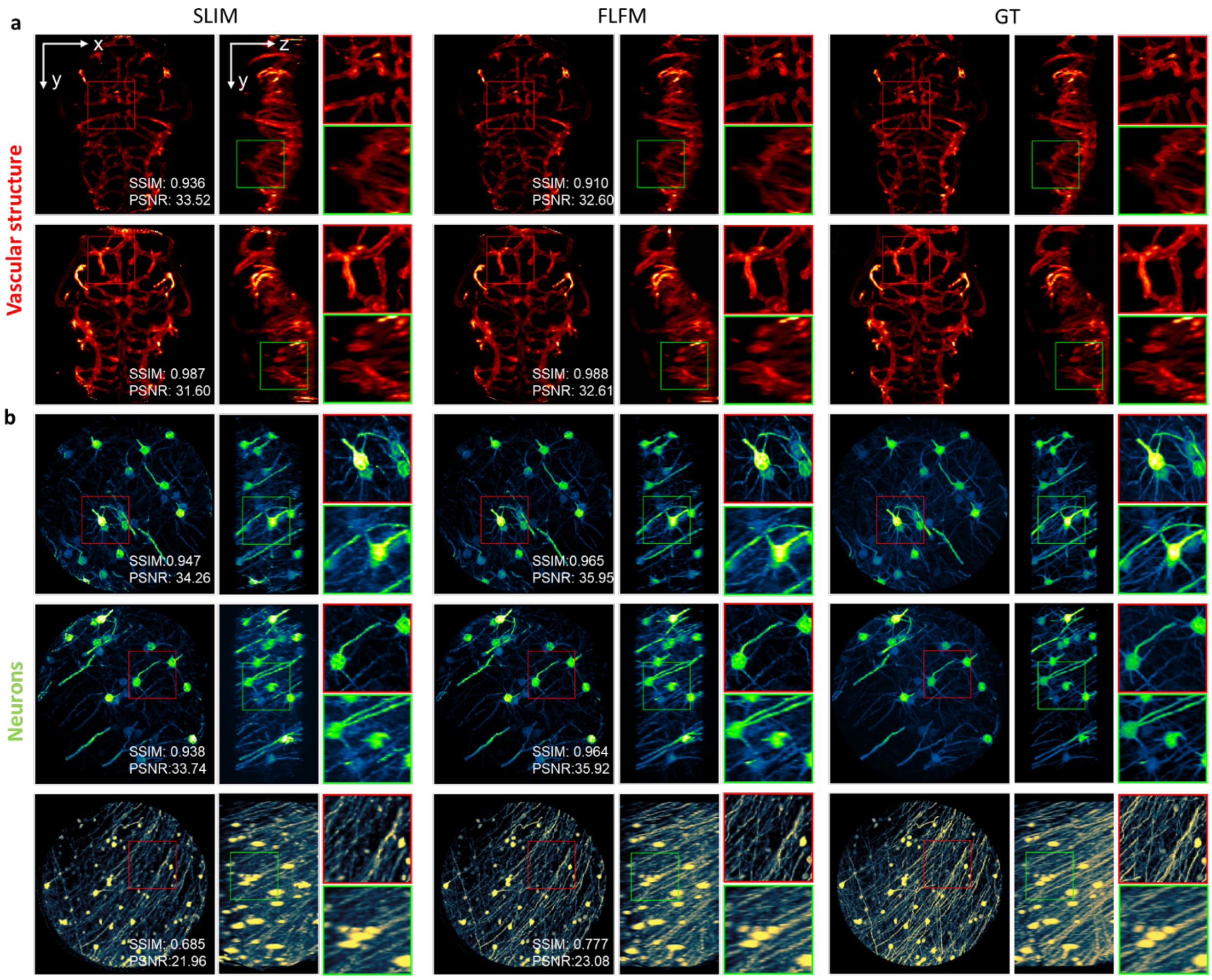


**Supplementary Note 2 Fig. 2. Comparisons of image quality between SLIM and FLFM in real biological sample simulations. a.**3D Maximum intensity projection (MIP) of the vascular structure in the brain of a transgenic zebrafish, *Tg(kdrl:HRAS-mCherry)*. **b.** 3D MIP of *Thy1-eGFP* (1st and 2nd rows) and *Thy1-eYFP* (3rd row) mouse neurons. The red (green) box highlights a zoomed-in view of the x-y (y-z) plane. Simulation adds Poisson noise to raw measurements. Sample sparseness is chosen to increase from top to bottom. Image reconstruction quality for SLIM and FLFM was compared across these samples using PSNR (Peak Signal-to-Noise Ratio) and SSIM (Structural Similarity Index Measure) as evaluation metrics.

### 2.**Number of sub-apertures and vertical scaling ratio**

In FLFM, the number of sub-apertures is associated with first-order parameters such as the NA of each sub-aperture, the FOV, the fill factor of the lenslet array (light efficiency), etc. In addition to these parameters, it determines the illness of reconstruction in SLIM, since fewer sub-aperture images make a higher compression ratio. Meanwhile, the vertical scaling ratio is defined by the demagnification of anamorphic relay. A smaller ratio allows a smaller sensor ROI and higher framerate but imposes higher vulnerability to noise and signal complexity.

The simulation in **Supplementary Note 2 Fig. 3** performs various scaling ratios with fixed number of sub-apertures (29). The ratio ranged from 0.04 to 1 and we tested on real zebrafish vascular structures and mouse brain neuron datasets. As the scaling ratio was reduced to extremely low values (e.g., 0.04 or 0.1), SLIM can still reconstruct signals across the entire FOV, but we observed image degradation, including increased artifacts and loss of fine details.

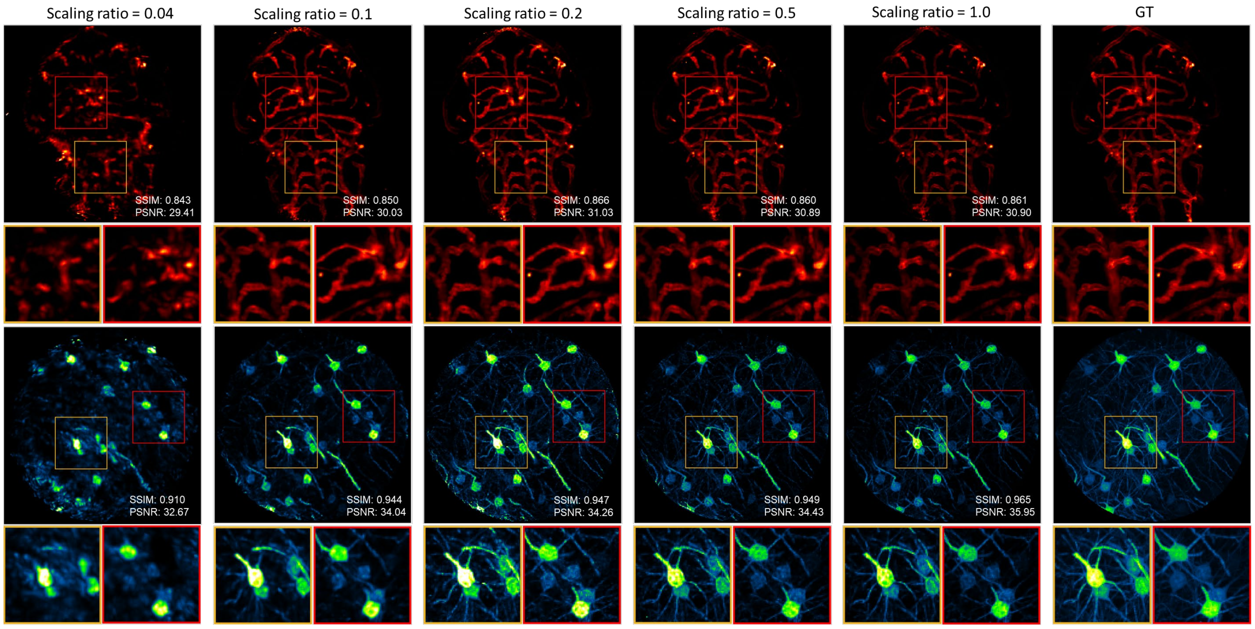


**Supplementary Note 2 Fig. 3. Comparisons of image quality of SLIM under different scaling ratio configurations**. Simulation adds Poisson noise. x-y MIPs are shown with zoom-in regions in red and yellow boxes. The reconstruction quality of SLIM and FLFM across different samples was compared using PSNR (Peak Signal-to-Noise Ratio) and SSIM (Structural Similarity Index Measure) as evaluation metrics.

Similarly, we controlled the number of sub-apertures (7, 19 and 29) but fixed the scaling ratio to 0.2. Simulations were performed on real zebrafish vascular structures and mouse brain neuron datasets (**Supplementary Note 2 Fig. 4**). As the number of sub-apertures decreased, we observed a degradation in image quality, manifesting as increased artifacts and a loss of detail. This decline occurs because fewer pixels are available for reconstruction, making the inverse problem more ill-posed. Here we changed the number of sub-apertures without adjusting lenslet size accordingly. Therefore, fewer sub-apertures also reduce light efficiency, compounding the negative effects on image quality.

Both the number of aperture and scaling ratio simulations reflect the inherent trade-off in the SLIM system. We experimentally demonstrated a scaling ratio of 0.2 and 29 apertures, enables kilohertz volumetric rate while maintaining reasonable robustness in various biological applications. However, the optimal SLIM configuration should consider specific sample structure, the targeted optical performance and framerate.

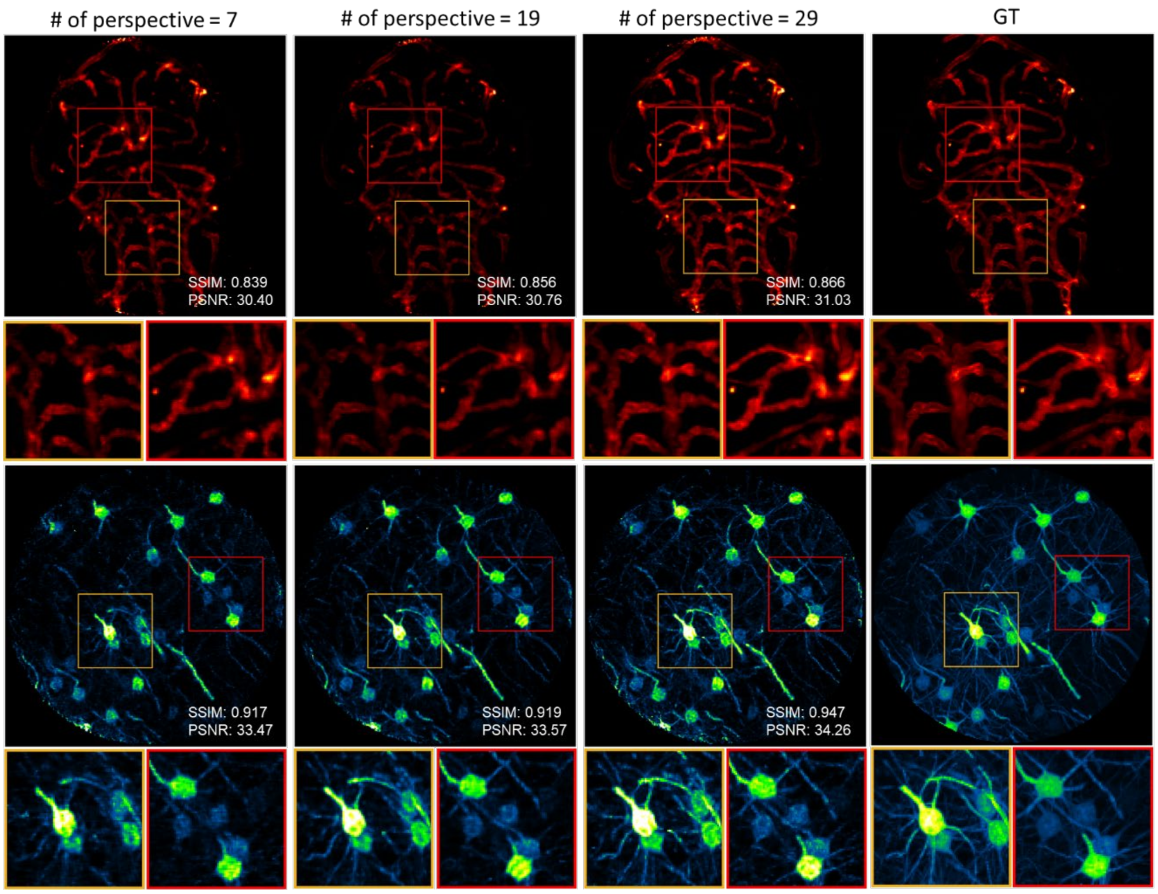


**Supplementary Note 2 Fig. 4. Comparisons of image quality of SLIM under different numbers of sub-apertures configurations**. Simulations add Poisson noise. x-y MIPs are shown with zoom-in regions in red and yellow boxes. The reconstruction quality of SLIM under different numbers of sub-apertures was compared using PSNR (Peak Signal-to-Noise Ratio) and SSIM (Structural Similarity Index Measure) as evaluation metrics.

### Supplementary Note 3 Pseudocode of SLIM reconstruction algorithm

**Algorithm** SLIM reconstruction using Richardson Lucy deconvolution

$s,\ \theta(N_v),\ PSF(N_x, sN_y, N_z, N_v),\ im(N_x, sN_y, N_v)$
    ▻ Input squeezing factor $s$, rotation angle $\theta$ , normalized PSF, and measured light field views $im$
$vol^0(N_x, N_y, N_z) = vol_{init}$ ▻ Initialize volume
**for** iterations **do**
    $\overline{im}$ = FP($vol^k$) ▻ Forward project the current volume estimation
    $vol^{k+1}$=$vol^k$ · BP($im/\overline{im}$) ▻ Backward project the error and update volume
**end for**

**function** FP($vol$) ▻ Forward project the volume of dimensions $N_x, N_y, N_z$
    **Initialize** $im = zeros(N_x, s \cdot N_y, N_v)$
    **for** each $v$ in $N_v$ **do**
        Apply in-plane rotation ($\theta(v)$) to entire $vol$
        Resize image at scale $s$ in direction $y$ for entire $vol$
        Slice-by-slice convolution between resized $vol$ and $PSF(:,:,:,v)$
        $im(:,:,v)$ = Sum along depth of the convolution result
    **end for**
    **return** $im$
**end function**

**function** BP($im$) ▻ Backward project the light field views of dimensions $N_x, s \cdot N_y, N_v$
    **Initialize** $vol = zeros(N_x, N_y, N_z)$
    **for** each $v$ and $d$ in $N_v$ and $N_z$ **do**
        $PSF^T(:,:,d,v)$ = rotate $PSF(:,:,d,v)$ at 180 degrees
    **end for**
    **for** each $d$ in $N_z$ **do**
        View-by-view convolution between $im$ and $PSF^T(:,:,d,:)$
        Resize convoluted views at scale $1/s$ in direction y
        Apply in-plane rotation to each view at corresponding angle $-\theta(v)$
        $vol(:,:,d)$ = Sum along view of rotated results
    **end for**
    **return** $vol$
**end function**

### Supplementary Table 1. Degraded performance in ultra-fast cameras/camera modes

| Camera | Quantum efficiency | Readout noise($e^-$) | Full well capacity($e^-$) | Bitdepth (bit) | Frame format (px) | FPS | On-board RAM (Max. recording time) |
|---|---|---|---|---|---|---|---|
| Hamamatsu ORCA Flash4.0 V3 | 82% | 1.6 | 30000 | 16 | 2048×2048 | 100 | |
| **Teledyne Kinetix (Dynamic range mode)** | **96%** | **1.6** | **15000** | 16 | 3200×3200 | 83 | |
| Teledyne Kinetix (Speed mode) | 96% | 2.0 | 200 | 8 | 3200×3200 | 500 | |
| Lambert HiCAM Fluo 2000[8,9] | 50% | 24 | | 8 | 1920×1080 | 2,000 | |
| Gpixel GSPRINT4502 (cropped FOV)[9] | 60% | 7 | 7400 | 10 | 1280×512 | 3,900 | |
| Phantom TMX 7510 | 77.6% | 24.18 | 8736 | 12 | 1280×800 | 76,000 | 512 GB (4.4s) |

### Supplementary Table 2. Space-bandwidth product (SBP) and volume rate of representative 3D microscopes

| | Method | FOV(μm) | Resolution (x,y,z, μm) | Measurement (px) | Volume rate (vps) | SBP×volume rate* | Detector | Voltage imaging |
|---|---|---|---|---|---|---|---|---|
| Compressive Fourier light field microscopy | **SLIM (this work)** | Ø550×300 | 3.6×3.6×6 | 3200×200 | 1000** | **7.3×10$^{9}$** | Teledyne Kinetix (16-bit dynamic range mode), sCMOS | Leech ganglion ex *vivo*, Mouse brain *in vivo* |
| | **Fourier DiffuserSco-pe**[10] | 1000×1000 ×280 | 3×3×4 | 4.2×10$^{6}$ | 25 | 1.56×10$^{9}$ | Andor Zyla 4.2, sCMOS | N.A. |
| | **Miniscope-3D**[11] | 900×700 ×390 | 2.76×2.76 ×15 | 0.3×10$^{6}$ | 40 | 6.88×10$^{8}$ | Ximea MU9PM-MH, CMOS | N.A. |
| Event camera | **Event LFM**[12] | 130×130 ×200 | 3.9×3.9×21 | 1280×720 | 1000 | 8.5×10$^{7}$ | EVK4, Prophesee, IMX636 sensor, Event Camera | N.A. |
| Deep learning enhancement | **Virtual Scanning LFM**[13] | 210×210×18 | 0.23×0.23 ×0.42 | 2048×2048 | 12 (63×/1.4 Oil) | 3.43×10$^{9}$ | Andor Zyla 4.2 Plus PCIE, sCMOS | N.A. |
| | | 260×260 ×100 | | 2000×2000 | 500 (25×/1.05 Water) | | Teledyne Kinetix (8-bit speed mode), sCMOS | *Drosophila* brain, *in vivo* |
| Multiple cameras | **CALM**[14] | 2650×2650 ×300 | 7.68×7.68 ×8.8 | 25×1024×768 | 30 | 9.74×10$^{8}$ | *****25×**PointGray Flea2-08S2C-C RGB, CMOS | N.A. |
| | **Wang et al.**[9] | 930×370 ×170 | 1.46×1.46 ×11.7 | 2×1280×256×30 | 200.8 | 3.77×10$^{9}$ | *****2×**Gpixel GSPRINT4521/10/02 | Zebrafish brain *in vivo* |
| Multi-plane scanning | **MuZIC**[15] | 150×150×45 (4 planes) | 2.6×1.9×12 (interplane spacing) | | 916 | 6.68×10$^{7}$ | Hamamatsu S14420-1550MG (SiPM) | Mouse brain *ex vivo* and *in vivo* |
| Axially swept light sheet | **SIFT**[16] | 4200 ×3300×5 500 | 0.97×0.97 ×0.97 | | 5.63×10$^{-5}$ (4.93h acquisition time) | 3.76×10$^{7}$ | Hamamatsu Orca Flash 4.0, sCMOS | N.A. |

*SBP is defined by $SBP = FOV/(d_x d_y d_z) * 8$, where $FOV$ is three-dimensional, $d$ is the spatial resolution, 8 accounts for the Nyquist sampling theorem. This definition refers to Lin et al.[14]

**The camera supports 1326 vps in 16-bit and 7476 vps in 8-bit. We only experimentally demonstrated 1000 vps in 16-bit and 4800 vps in 8-bit in accommodation to sample signal SNR.

***The number of detectors in the array.

### Supplementary Table 3. List of components used in SLIM

| | | Component | Description | Manufacturer | Part number | Material |
|---|---|---|---|---|---|---|
| Selective volume illumination setup | SLIM | O1 | 20X Olympus XLUMPLFLN Objective, 1.00 NA | Olympus | N20X-PFH | |
| | | L1 | 180 mm focal length achromatic doublet | Thorlabs | AC508-180-A | |
| | | L2 | 200 mm focal length achromatic doublet | Thorlabs | AC508-200-A | |
| | | L3 | 250 mm focal length achromatic doublet | Thorlabs | ACT508-250-A | |
| | | CL1 | 250 mm focal length cylindrical achromatic doublet | Thorlabs | ACY254-250-A | |
| | | CL2 | 50 mm focal length cylindrical achromatic doublet | Thorlabs | ACY254-50-A | |
| | | Dove prism | 1.3mm aperture size customized dove prism | Changchun Sunday Optics | | H-K9L glass |
| | | Dove prism holder | Customized 3D printing holder | Protolabs | | Accura 7820 3D Printing Material |
| | | MLA | 36 mm focal length plano-convex lenslet array | fabricated in-house | | PMMA |
| | | Camera | sCMOS camera | Teledyne | Kinetix | |
| | | F1(GFP) | 525nm center wavelength bandpass filter | Chroma | ET525/50m | |
| | | F1(RFP) | 585nm center wavelength bandpass filter | Chroma | ET585/65m | |
| | Dual Scanning light sheet | 473nm laser | 500mW 473nm diode laser | CNI laser | MBL-FN-473-500mW | |
| | | 532nm laser | 300mW 532nm diode laser | CNI laser | MGL-III-532-300mW | |
| | | Beam splitter | 50:50 Non-Polarizing Beamsplitter Cube | Thorlabs | BS013 | |
| | | CL3 | 50 mm focal length cylindrical lens | Thorlabs | LJ1695RM-A | |
| | | CL4 | 50 mm focal length cylindrical lens | Thorlabs | LJ1695RM-A | |
| | | Knife-edge mirror | Knife-edge right-angle prism | Thorlabs | MRAK25-G01 | |
| | | L4 | 150 mm focal length achromatic doublet | Thorlabs | AC254-150-A | |
| | | Galvo mirror | single axis scanning Galvo mirror | Thorlabs | GVS011 | |
| | | L5 | 150 mm focal length achromatic doublet | Thorlabs | AC508-150-A | |
| | | L6 | 180 mm focal length achromatic doublet | Thorlabs | AC508-150-A | |
| | | O2 | 4X Olympus Plan Fluorite Objective, 0.13 NA | Olympus | RMS4X-PF | |
| | LED | Blue LED | ultra-low-noise blue LED | Prizmatix | UHP-T-470SR | |
| | | Excitation filter | 470nm center wavelength bandpass filter | Chroma | ET470/40X | |
| | | L7 | 50 mm focal length achromatic doublet | Thorlabs | AC254-050-A | |
| | | Slit | adjustable mechanical slit | Thorlabs | VA100 | |
| | | L8 | 80 mm focal length achromatic doublet | Thorlabs | AC508-080-A | |
| | | O2 | 4X Olympus Plan Fluorite Objective, 0.13 NA | Olympus | RMS4X-PF | |
| Widefield illumination setup | SLIM | O3 | 16X Nikon CFI LWD Plan Fluorite Objective, 0.80 NA | Nikon | N16XLWD-PF | |
| | | L9 | 150 mm focal length achromatic doublet | Thorlabs | AC508-150-A | |
| | | L10 | 150 mm focal length achromatic doublet | Thorlabs | AC508-150-A | |
| | | L11 | 250 mm focal length achromatic doublet | Thorlabs | ACT508-250-A | |
| | | CL5 | 250 mm focal length cylindrical achromatic doublet | Thorlabs | ACY254-250-A | |
| | | CL6 | 50 mm focal length cylindrical achromatic doublet | Thorlabs | ACY254-50-A | |
| | | Dove prism | 1.3mm aperture size customized dove prism | Changchun Sunday Optics | | H-K9L glass |
| | | Dove prism holder | Customized 3D printing holder | Protolabs | | Accura 7820 3D Printing Material |
| | | MLA | 36 mm focal length plano-convex lenslet array | fabricated in-house | | PMMA |
| | | Camera | sCMOS camera | Teledyne | Kinetix | |
| | | F2 | 525nm center wavelength bandpass filter | Chroma | ET525/50m | |
| | LED | Blue LED | Ultra-low-noise blue LED | Prizmatix | UHP-T-470SR | |
| | | F3 | 470nm center wavelength bandpass filter | Chroma | ET470/40X | |
| | | F4 | GFP Dichroic Filter | Thorlabs | MD498 | |
| | | L12 | 150 mm focal length achromatic doublet | Thorlabs | AC254-150-A | |

### Supplementary Table 4. Acquisition parameters for imaging experiments

| | Figure Number | Sample | Illumination Source | Galvo Mirror Frequency (Hz) | Sensor ROI Size (px) | Sensor Readout Mode | Sensor Exposure Time (µs) | Sensor Frame Rate (fps) | Volume Rate (vps) | Recording Time (s) | Number of sub-apertures | Reconstruction Resolution (px) | Reconstruction Pixel Size (µm) | FOV(µm) | Illumination Power ( mW/mm2) |
|---|---|---|---|---|---|---|---|---|---|---|---|---|---|---|---|
| Selective volume illumination setup | Fig. 1e,f | Fluorescent beads (F13081, ThermoFisher) | LED (470 nm) | Not applicable | 320×3200 | 16-bit (Dynamic range mode) | 100000 | Not applicable | Not applicable | Not applicable | 29 | 305×305×101 | 1.81×1.81×4 | Ø550×400 | |
| | Fig. 2a-c | Embryonic zebrafish *Tg(gata1a:dsRed)* @ 3dpf | Scanning light sheet (532 nm) | 1000 | 200×3200 | 16-bit (Dynamic range mode) | 990 | 1000 | 1000 | 4 | 19 | 305×305×51 | 1.81×1.81×6.16 | Ø550×308 | |
| | Fig. 2d-e | Embryonic zebrafish *Tg(gata1a:dsRed)* @ 3dpf | Scanning light sheet (532 nm) | 1000 | 200×3200 | 16-bit (Dynamic range mode) | 990 | 1000 | 1000 | 4 | 19 | 305×305×151 | 1.81×1.81×4 | Ø550×600 | |
| | Fig. 3c,f, S7 | Medicinal leech | LED (470 nm) | Not applicable | 320×3200 | 16-bit (Dynamic range mode) | 1200 | 800 | 800 | 12 | 29 | 295×295×76 | 1.81×1.81×4 | Ø532×300 | |
| | Fig. 5b | Fluorescent beads (F13082, ThermoFisher) | Scanning dual-light sheet (532 nm) | 100 | 320×3200 | 8-bit (Speed mode) | 240 | 4000 | 100 | Not applicable | 29 | 295×295×78 | 1.81×1.81×3.9 | Ø532×300 | |
| | Fig. 5c | Embryonic zebrafish *Tg* (flk:mCherry) @ 3dpf | Scanning dual-light sheet (532 nm) | 100 | 320×3200 | 8-bit (Speed mode) | 240 | 4000 | 100 | 3 | 29 | 295×295×78 | 1.81×1.81×3.9 | Ø532×300 | |
| | Fig. 5d,e,f | Embryonic zebrafish *Tg* (cmlc:GFP) @ 3dpf | Scanning dual-light sheet (473 nm) | 300 | 200×3200 | 8-bit (Speed mode) | 200 | 4800 | 300 | 3 | 19 | 150×150×30 | 1.81×1.81×6 | Ø270×174 | |
| | Fig. S14 | Vibrio cholerae bacteria | Scanning light sheet (473nm) | 200 | 320×3200 | 16-bit (Dynamic range mode) | 4800 | 200 | 200 | 5 | 29 | 295×295×76 | 1.81×1.81×4 | Ø532×300 | |
| Widefield illumination setup | Fig. 4, Ext. Fig. 2, Fig. S12, S13 | Wild type mice with genetically encoded voltage indicator (GEVI) pAce (interneurons) | LED (470 nm) | Not applicable | 330×2400 | 16-bit (Dynamic range mode) | 1230 | 800 | 800 | 180 | 29 | 305×305×75 | 2.26×2.26×8 | Ø686×592 | 40 |
| | Fig. S14, S15 | Wild type mice with genetically encoded voltage indicator (GEVI) pAce (interneurons) | LED (470 nm) | Not applicable | 330×2400 | 16-bit (Dynamic range mode) | 1230 | 800 | 800 | 180 | 29 | 305×305×75 | 2.26×2.26×8 | Ø686×592 | 36 |
| | Ext. Fig 1 | Wild type mice with genetically encoded voltage indicator (GEVI) pAce (pyramidal neurons) | LED (470 nm) | Not applicable | 330×2400 | 16-bit (Dynamic range mode) | 1230 | 800 | 800 | 180 | 29 | 305×305×75 | 2.26×2.26×8 | Ø686×592 | 25 |
| | Fig. S11 | Fluorescent beads (F13081, ThermoFisher) | LED (470 nm) | Not applicable | 330×2400 | 16-bit (Dynamic range mode) | 1230 | Not applicable | Not applicable | Not applicable | 29 | 305×305×151 | 2.26×2.26×8 | Ø686×1200 | |

### References

1. Guo, C. *et al.* Fourier light-field microscopy. *Opt. Express, OE* **27**, 25573–25594 (2019).
2. Vladimirov, N. *et al.* Light-sheet functional imaging in fictively behaving zebrafish. *Nat Methods* **11**, 883–884 (2014).
3. Cong, L. *et al.* Rapid whole brain imaging of neural activity in freely behaving larval zebrafish (Danio rerio). *eLife* **6**, e28158 (2017).
4. Kugler, E. C. *et al.* Zebrafish vascular quantification: a tool for quantification of three-dimensional zebrafish cerebrovascular architecture by automated image analysis. *Development* **149**, dev199720 (2022).
5. Dean, K. M. *et al.* Isotropic Imaging Across Spatial Scales with Axially Swept Light-Sheet Microscopy. *Nat Protoc* **17**, 2025–2053 (2022).
6. Park, H. *et al.* Deep learning enables reference-free isotropic super-resolution for volumetric fluorescence microscopy. *Nat Commun* **13**, 3297 (2022).
7. Yoon, Y.-G. *et al.* Sparse decomposition light-field microscopy for high speed imaging of neuronal activity. *Optica, OPTICA* **7**, 1457–1468 (2020).
8. Mandracchia, B. *et al.* High-speed optical imaging with sCMOS pixel reassignment. *Nat Commun* **15**, 4598 (2024).
9. Wang, Z. *et al.* Imaging the voltage of neurons distributed across entire brains of larval zebrafish. 2023.12.15.571964 Preprint at https://doi.org/10.1101/2023.12.15.571964 (2023).
10. Liu, F. L., Kuo, G., Antipa, N., Yanny, K. & Waller, L. Fourier DiffuserScope: single-shot 3D Fourier light field microscopy with a diffuser. *Opt. Express, OE* **28**, 28969–28986 (2020).

11. Yanny, K. *et al.* Miniscope3D: optimized single-shot miniature 3D fluorescence microscopy. *Light Sci Appl* **9**, 171 (2020).

12. Guo, R. *et al.* EventLFM: event camera integrated Fourier light field microscopy for ultrafast 3D imaging. *Light Sci Appl* **13**, 144 (2024).

13. Lu, Z. *et al.* Virtual-scanning light-field microscopy for robust snapshot high-resolution volumetric imaging. *Nat Methods* **20**, 735–746 (2023).

14. Lin, X., Wu, J., Zheng, G. & Dai, Q. Camera array based light field microscopy. *Biomed. Opt. Express, BOE* **6**, 3179–3189 (2015).

15. Weber, T. D., Moya, M. V., Kılıç, K., Mertz, J. & Economo, M. N. High-speed multiplane confocal microscopy for voltage imaging in densely labeled neuronal populations. *Nat Neurosci* **26**, 1642–1650 (2023).

16. Prince, M. N. H. *et al.* Signal improved ultra-fast light-sheet microscope for large tissue imaging. *Commun Eng* **3**, 1–13 (2024).
